# Dimerization of kringle 1 domain from hepatocyte growth factor/scatter factor provides a potent minimal MET receptor agonist

**DOI:** 10.1101/2020.07.20.212654

**Authors:** Giovanni de Nola, Bérénice Leclercq, Alexandra Mougel, Solenne Taront, Claire Simonneau, Federico Forneris, Eric Adriaenssens, Hervé Drobecq, Luisa Iamele, Laurent Dubuquoy, Oleg Melnyk, Ermanno Gherardi, Hugo de Jonge, Jérôme Vicogne

## Abstract

Hepatocyte Growth Factor/Scatter Factor (HGF/SF) and its cognate receptor MET play several essential roles in embryogenesis and regeneration in post-natal life of epithelial organs such as liver, kidney, lung, and pancreas, prompting a strong interest in harnessing HGF/SF-MET signalling for regeneration of epithelial organs after acute or chronic damage. The limited stability and tissue diffusion of native HGF/SF, however, which reflect the tightly controlled, local mechanism of action of the morphogen, have led to a major search for HGF/SF mimics for therapy. In this work, we describe the rational design, production and characterisation of K1K1, a novel minimal MET agonist consisting of two copies of the kringle 1 domain of HGF/SF placed in tandem. K1K1 is highly stable and displays biological activities equivalent or superior to native HGF/SF in a variety of *in vitro* assay systems and in a mouse model of liver disease. These data suggest that this engineered ligand may find wide applications in acute and chronic diseases of the liver and other epithelial organs dependent on MET activation.

## Introduction

Hepatocyte Growth Factor/Scatter Factor (HGF/SF) was simultaneously and independently discovered as a hepatotropic factor in the serum of hepatectomized rats (Nakamura *et al*, 1984) and as a scatter factor (SF) inducing morphological changes and epithelial cell migration in medium conditioned by human embryo fibroblasts (Stoker & Perryman, 1985; Stoker *et al*, 1987). After the discovery of its cognate receptor, the tyrosine kinase receptor MET (Bottaro *et al*, 1991), important studies using targeted gene disruption or blocking antibodies demonstrated embryonic developmental defects in placenta (Uehara *et al*, 1995), liver (Schmidt *et al*, 1995), skeletal muscles of limb and diaphragm (Bladt *et al*, 1995), and kidney (Santos *et al*, 1994; Woolf *et al*, 1995). Collectively, these studies demonstrate an early and wide involvement of HGF/SF-MET signalling in organ development.

In post-natal life, the plasma level of HGF/SF rises immediately after injury of several organs such as liver, kidney, and heart (Lindroos *et al*, 1991; Nakamura *et al*, 2000; Zhu *et al*, 2000; Matsumoto & Nakamura, 1997). HGF/SF is essential for the regeneration of kidney (Zhou *et al*, 2013), liver (Borowiak *et al*, 2004), and skin (Chmielowiec *et al*, 2007) and, importantly, treatments with recombinant HGF/SF prevents fibrosis after experimental injury in kidney (Liu, 2004) and lung (Mizuno *et al*, 2005) providing a strong rationale for harnessing HGF/SF-MET signalling for regenerative medicine.

The mature, biologically active species of HGF/SF is a two-chain (α/β), disulphide-linked protein structurally related to the blood proteinase plasminogen. The α-chain (69 kDa) comprises an N-terminal (N) domain homologous to the plasminogen activation peptide and four kringle (K) domains and contains the high affinity-binding site for MET. The β-chain (34 kDa) is homologous to the catalytic domain of serine proteinases (SPH) (Donate *et al*, 1994); it is devoid of enzymatic activity but contains a secondary binding site for MET. During tissue remodelling and at sites of tissue damage, the activation of the clotting/fibrinolytic pathways promotes the cleavage of the single chain pro-HGF/SF into mature two-chain HGF/SF leading to MET activation. In this way, HGF/SF-MET signalling promotes cell survival, division, migration and a complex morphogenetic program underlying tissue regeneration (Miyazawa *et al*, 1994, 1996).

The discovery of two alternative HGF/SF splice variants, NK1 and NK2 (Miyazawa *et al*, 1991; Chan *et al*, 1991; Cioce *et al*, 1996) enabled further progress toward the identification of regions important for receptor binding and activation (Hartmann *et al*, 1992; Lokker *et al*, 1992). NK1, a partial receptor agonist containing the N-domain and first kringle domain (K1) is monomeric in solution but assembles as a head-to-tail non-covalent dimer in crystal structures offering an attractive bivalent receptor-activating configuration (Chirgadze *et al*, 1999; Ultsch *et al*, 1998). Heparin and heparan sulphate proteoglycans cause dimerization of NK1 in solution (Schwall *et al*, 1996) and are essential for the partial agonistic activity of NK1 but not of HGF/SF (Zioncheck *et al*, 1995; Sakata *et al*, 1997; Schwall *et al*, 1996) and, on the strength of the available biochemical data and partial crystal structures, alternative models of HGF/SF-MET activation were proposed involving with 1:2 or 2:2 stoichiometries (Fig 1A) (Chirgadze *et al*, 1998; Donate *et al*, 1994; Miller & Leonard, 1998).

**Figure 1.**
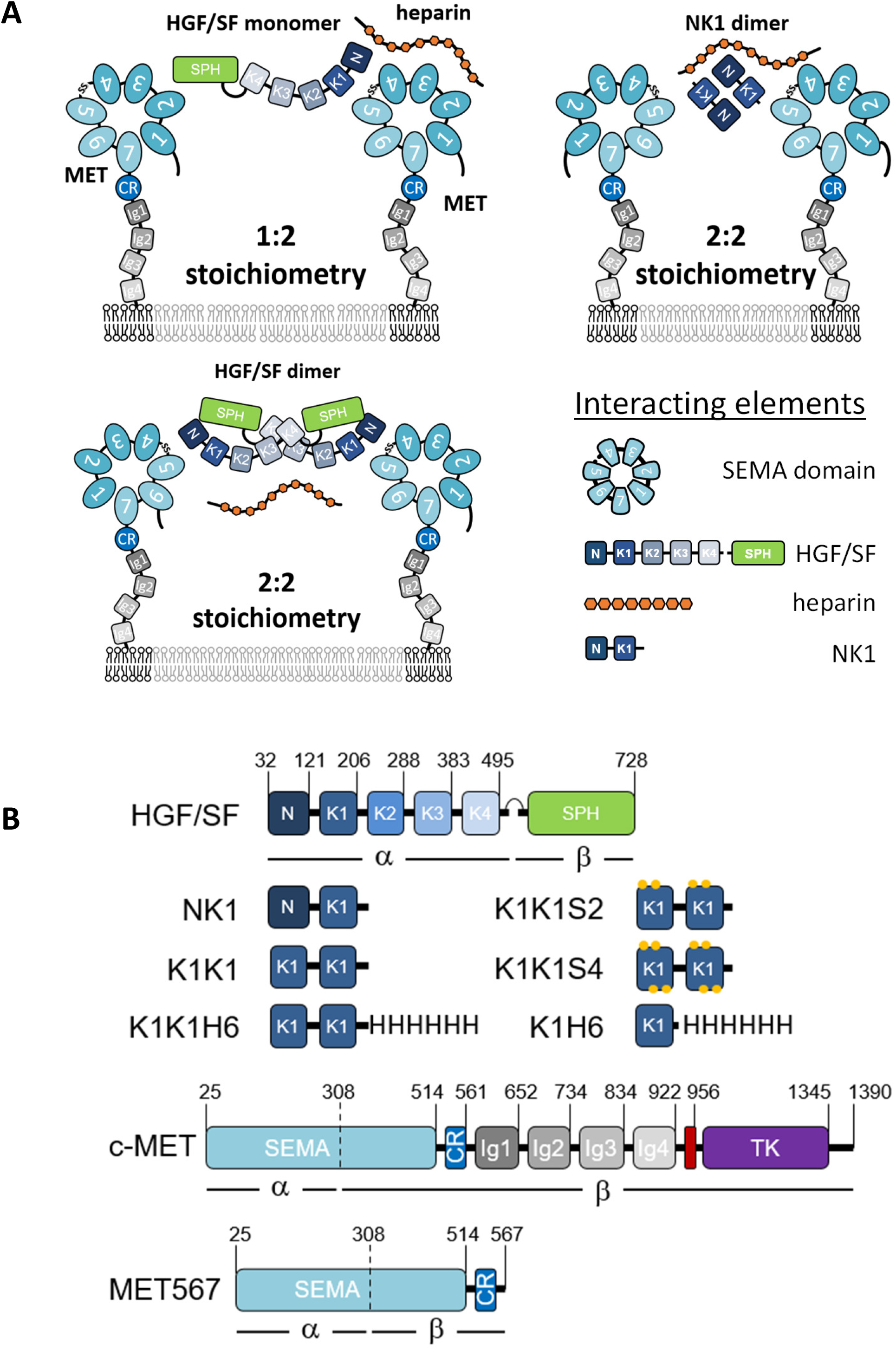
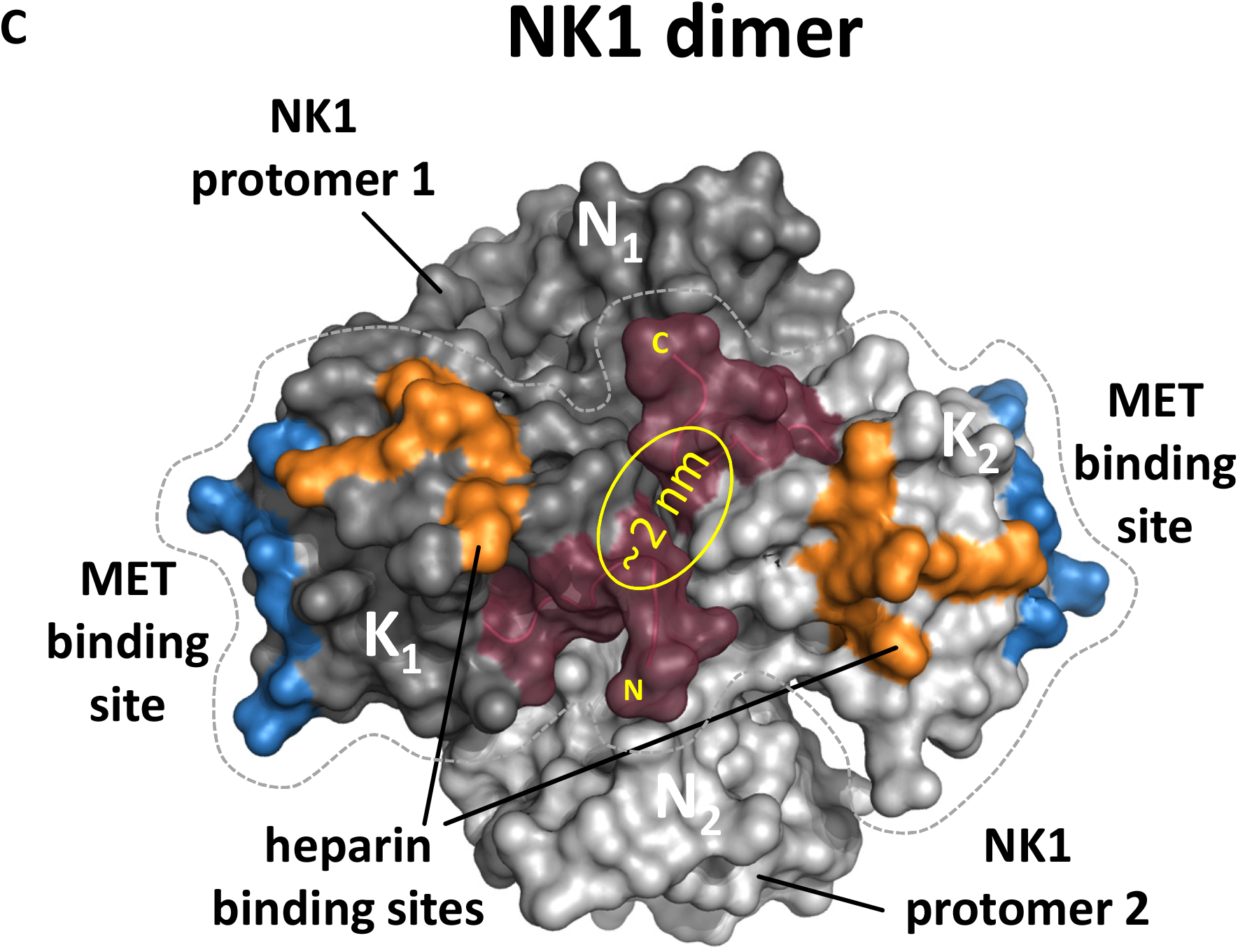
Potential receptor binding and activation modes, domain architecture of MET agonists, and surface representation of NK1 crystal structure. (A) Possible receptor binding modes and stoichiometries for HGF/SF and NK1 in the presence of heparin. (B) Schematic representation of HGF/SF, NK1, K1K1, K1K1 variants, K1H6, full length c-MET, and the MET567 fragment. Individual domains (boxes) with positions of domain boundaries indicated above. CR: cysteine rich, Ig: immunoglobulin-like, SPH: serine protease homology, TK: tyrosine kinase. The transmembrane domain is indicated in red. (C) Surface representation of the NK1 dimer with the two protomers shown in light and dark grey with residues involved in MET binding and heparin binding shown in blue and orange, respectively. The centrally placed N- and C-terminus that were connected by a linker are shown in Bordeaux.

A recent cryo-electron-microscopy (EM) structure of HGF/SF in complex with the extracellular domain of MET (MET927) fused to a dimerization motif has offered new insights into the binding mechanisms (Uchikawa *et al*, 2021). This structure unambiguously shows a single HGF/SF molecule bridging two MET receptors by engaging both the NK1 and SPH binding sites with a second HGF/SF molecule and heparin stabilising the interaction. In contrast, a second structure of the NK1-MET927 complex confirmed the presence of the complex with 2:2 stoichiometry postulated on the basis of the early crystal structures of NK1. This structure validates independent studies on the N and K1 domain of HGF/SF produced in yeast (Holmes *et al*, 2007) or by total chemical synthesis (Raibaut *et al*, 2013; Simonneau *et al*, 2015) suggesting K1 domain as the main or sole domain of HGF/SF responsible for the primary, high-affinity interaction between NK1 or HGF/SF and the SEMA domain of MET.

On the strength of the structural, biochemical and mutagenesis data available, we rationalised that a covalent dimer containing a tandem repeat of the K1 domain of HGF/SF could constitute a minimal and stable MET agonist able to diffuse effectively across tissue boundaries owing to the absence of the N-terminal domain of HGF/SF responsible for high-affinity binding to heparan sulfate proteoglycans (HSPG). Here we report the development of this new MET ligand which we name K1K1. We describe its structure and demonstrate that K1K1 is stable and displays biological activity, equivalent or superior compared to HGF/SF, in both *in vitro* and *in vivo* models. This new MET ligand may constitute a major step towards the prospect of harnessing MET signalling for the therapy of acute and chronic degenerative diseases of major epithelial organs.

## Results

### Rational for the design of K1K1

The molecular architecture of full MET receptor, its extracellular fragment and the various agonists used in this study are presented figure 1B. The concept of connecting two HGF/SF kringle 1 domains in tandem for the creation of the potent MET agonists described in this work comes from the initial observation that NK1 monomers form dimers upon crystallisation providing two MET receptor binding sites at opposite ends (Ultsch *et al*, 1998; Chirgadze *et al*, 1999). Within the structure of the NK1 dimer, the N- and C-termini of both kringle domains are all located in close proximity at the centre of the dimer structure, suggesting that a crosslinker could be made to link two kringle domains for mimicking the arrangement of K1 domains in NK1 dimeric complex. The recreation of such a configuration was first achieved using the streptavidin molecular platform combined with C-terminally biotinylatined synthetic kringle 1 (K1B) (Simonneau *et al*, 2015). Oligomerisation would place the K1B C-termini close together (~ 2 nm) around the streptavidin core and project the K1-MET binding site located around residue Glu159 away from it, mimicking the arrangement of K1 domains in NK1 crystal structures (Fig. 1C). The remarkable potency of streptavidin-K1B complex confirmed our assumption that multimerisation would lead to strong receptor activation and thus completed earlier observations for the location of the high-affinity receptor binding site in the first kringle domain (Lokker *et al*, 1994; Chirgadze *et al*, 1999; Gherardi *et al*, 2006). As potential heterogeneity of the streptavidin-K1B complex as well as the streptavidin moiety itself precluded its development as a therapeutic and would hamper structural studies, we hoped that strong agonistic activity could be achieved as well by associating two K1 domains, in tandem and covalently (Fig. 1C). Importantly, the activity of a such a covalent K1 dimer would no longer rely on a protein dimerization or multimerization assembly as required for wild-type NK1 (Sakata *et al*, 1997; Schwall *et al*, 1996) and streptavidin-K1B.

In wild-type NK1, the distance between the last cysteine in one kringle and the first cysteine in the other kringle is roughly 1 nm, giving possibility to associate the two kringle 1 domains in tandem. An asset of such a design is to enable the production of the molecule by recombinant methods. Therefore, we designed a linker to bridge the distance between the C-terminus of the first K1 domain and the N-terminus of the second one. We chose for that purpose the four amino acid long linker, SEVE, which is naturally present between kringle 1 and kringle 2 (K2) in HGF/SF. This native linker showed no contacts with neighbouring kringle domains in wildtype NK2 reported by Tolbert and colleagues (PDB code 3HN4) (Tolbert *et al*, 2010) and in NK2 in complex with heparin (PDB code 3SP8) and was therefore believed to not constrain the relative orientation of the K1 domains. Furthermore, a poly-histidine tagged variant, designated K1K1H6, was produced to facilitate certain assays as well as a single kringle 1 domain poly-histidine tagged variant (K1H6) as a monovalent control. As earlier studies had revealed a secondary low-affinity heparin binding site in the kringle 1 domain affecting biological activity (Lietha *et al*, 2001), we designed two additional variants, K1K1S2 and K1K1S4. In these variants, we have introduced 2 or 4 reverse-charge mutations in each kringle (schematically shown in Fig. 1B) in order to test the implication of the basic residues naturally present at those positions and to evaluate their importance for the biological activity.

Upon expression in *Escherichia coli*, K1K1 was abundantly present in inclusion bodies. Expression at 18 °C and induction with a low IPTG concentration (0.1 mM) allowed the production and extraction of fully folded protein from “non-classical” inclusion bodies using a mild, non-denaturing arginine-based extraction method derived from the work of Jevševar and colleagues (Jevševar *et al*, 2005). Expression in these inclusion bodies offers an inexpensive and abundant source of recombinant protein with relatively few contaminants as opposed to soluble bacterial expression. The low-affinity heparin binding site in the kringle domain still allowed an effective single-step purification by heparin Sepharose^®^ affinity chromatography following arginine extraction (Fig. S1A). An additional size exclusion chromatography step was used to remove possible traces of aggregation to yield crystallography-grade material with UPLC-MS analysis confirming the purity and identity of the proteins as shown in Fig. S1B and S1C for K1K1 and K1K1H6 respectively. Figure S1D shows all purified recombinant proteins used in this study on Coomassie-stained gel under reducing conditions.

### Structure determination by x-ray diffraction and SAXS

Purified proteins were used for crystallization experiments using commercial sparse matrix screens and several conditions resulted in the growth of protein crystals at 17 °C within days. The collected datasets enabled the determination of the molecular structures at 1.8 Å resolution and 1.7 Å for K1K1 and K1K1H6 respectively. Both proteins crystallized in space group P 1 21 1 with K1K1 having two molecules per asymmetric unit and K1K1H6 having one.

Structurally, K1K1 and K1K1H6 are nearly identical with a backbone root mean square deviation (RMSD) of 0.8 or 1.8 Å (Fig. 2B), depending on which of the two molecules in the asymmetric unit of K1K1 is used for alignment with K1K1H6. Both proteins show an elongated “stretched-out” conformation with the MET binding sites exposed on opposite ends and the N- and C-terminus located at the centre. In K1K1H6, good electron density was also observed for the poly-histidine tag possibly due to stabilizing interactions between the histidine residues and residues of the N-terminal kringle domain.

**Figure 2.**
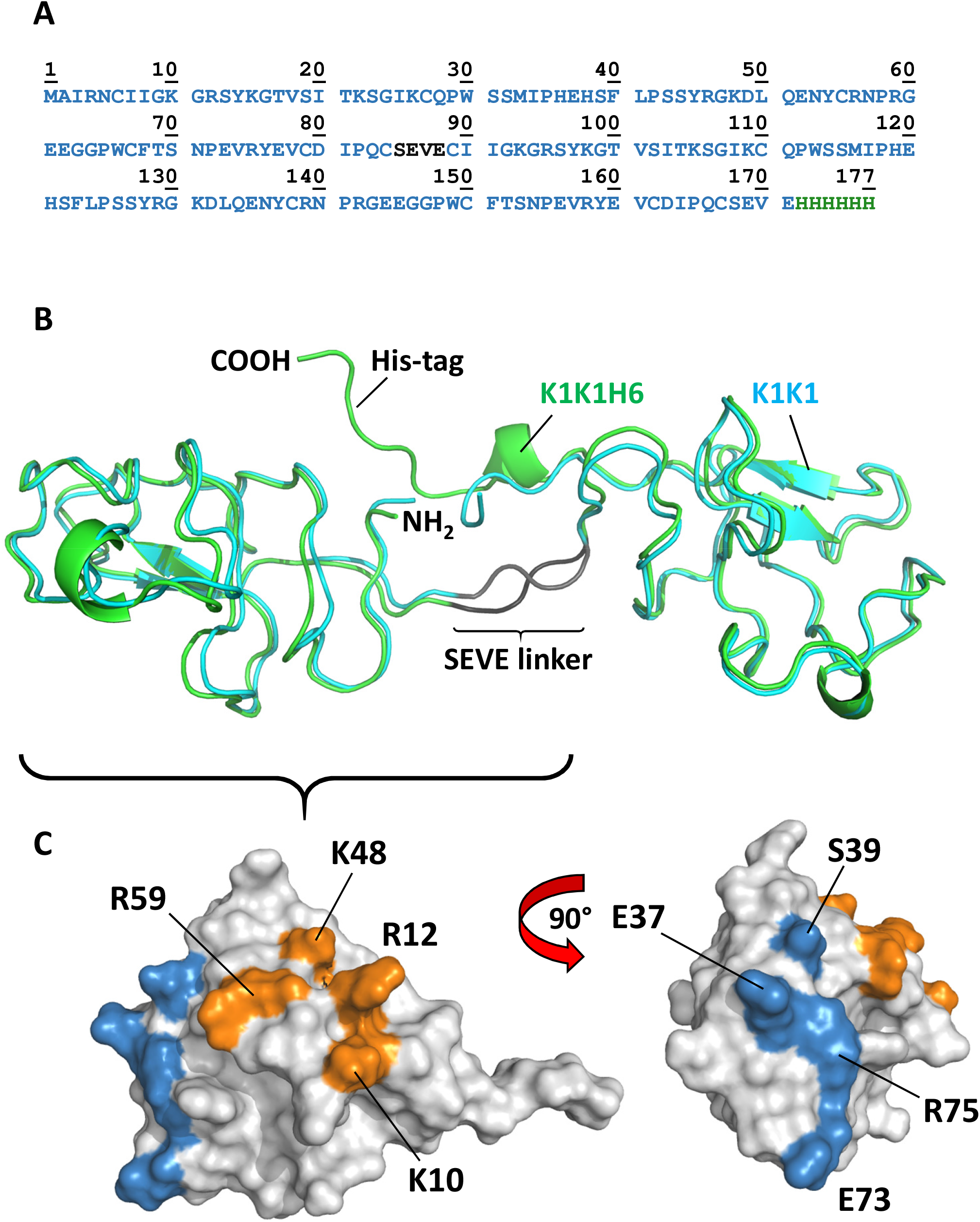
Amino acid sequence, overall structure, and location of binding sites. (A) Amino acid sequence of K1K1H6 with SEVE linker in dark grey and poly-histidine tag in green. (B) The crystal structure of the two molecules of K1K1 (cyan) and K1K1H6 (green) showing the straight conformation and the N-terminus (NH2) and C-terminus (COOH) located centrally in the linker region and a nearly identical overall structure with an RMSD ranging from 0.8 to 1.8 Å. The C-terminal poly-histidine tag of K1K1H6 is making contacts with residues in the N-terminal kringle domain. (C) Surface representations of single kringle domains of K1K1 showing the location of the residues involved in MET binding, as defined by Lokker et al., 1994 40, shown in blue. The more lateral position of the heparin binding site is shown in orange with the residues targeted by reverse charge mutation in K1K1S2, (K10E, R12E) and K1K1S4, (K10E, R12E, K48E, R59E) indicated.

Surprisingly, like the revelation of seeing the first structures of NK1 that presented a likely biologically relevant dimer, both crystal structures show a pseudo C2 symmetry around a central axis and present both receptor binding sites in nearly identical orientation on either side of the molecule (located around residue Glu37 in the first kringle and residues Glu120 in the second kringle, equivalent to NK1 residue Glu159). One can envision a binding mode in which two MET receptor monomers are brought in proximity by binding on opposite sides of K1K1, forming a 2:1 receptor activation complex. K1K1 therefore presents the most minimalistic peptide-based receptor agonist mimicking the binding and receptor activation mechanism proposed for wild-type NK1 but with the important difference that ligand dimerization is not required. This covalent “mimicry” is even more evident when comparing the structures of the NK1 dimer and K1K1. Alignment of the first kringle domain of K1K1 with one of the two kringle domains within the NK1 dimer places the second K1K1 kringle close to the position of the kringle domain of NK1 protomer 2 (Fig. S2A). However, the straightened conformation observed in the crystal structure of K1K1, “misaligns” the second K1K1 kringle domain through a rotation by 109.7 degrees and a translation of roughly 14 Å (Fig. S2B). No contacts between the two K1 domains or the kringle domains with the SEVE linker are observed. The linker is more stretched out compared to the two NK2 structures supporting our idea that it is highly flexible and therefore allows significant conformational freedom (Fig. S2C). This flexibility was confirmed with SAXS measurements of K1K1 in solution which generated an envelope that perfectly accommodates K1K1 volume-wise but only in a bent conformation (Fig. 3A). This is also evident from comparing the experimental scatter curve with the one generated based on the K1K1 crystal structure using CRYSOL indicating that the overall volume is the same but the distribution and therefore conformation differs significantly. Notably, the K1K1 solution SAXS envelope is compatible with the two kringle domains in the orientation found within the NK1 dimer (Fig. S2D). Excitingly, SAXS measurements of K1K1 in complex with MET567 gave a molecular mass estimation of 86.8 kDa and a calculated envelope that corresponds to a 1:1 complex with a “pan handle” extrusion which could accommodate K1K1 bound to the SEMA domain. Using a low-resolution crystallographic dataset of NK1 in complex with MET567 (unpublished, briefly described in Blaszczyk et al. (Blaszczyk *et al*, 2015)), we could generate a model based on the alignment of K1K1 with the crystal structure of the NK1-MET567 complex. The theoretical scatter curve matched the experimental scatter curve remarkably well (Chi^2^=3.6) with also in this model a slightly bent K1K1 conformation fitting best with the SAXS envelope. Moreover, the recently published cryo-EM structure of the NK1 dimer in complex with two SEMA domains (Uchikawa *et al*, 2021) perfectly accommodates our K1K1 SAXS envelope overlapping both NK1 kringle domains and supports the proposed model for receptor activation by K1K1 (Fig. 3C, Fig. S2E). Overall, we have generated compelling structural support for a minimal MET receptor agonist that comprises two covalently linked MET binding sites (see Supplementary Video 1). The structure of K1K1 and K1K1H6 are available from the PDB data bank with id code 7OCL and 7OCM respectively. The SAXS data is available from SASBDB database with id codes SASDLM9, SASDLN9, and SASDLP9.

**Figure 3.**
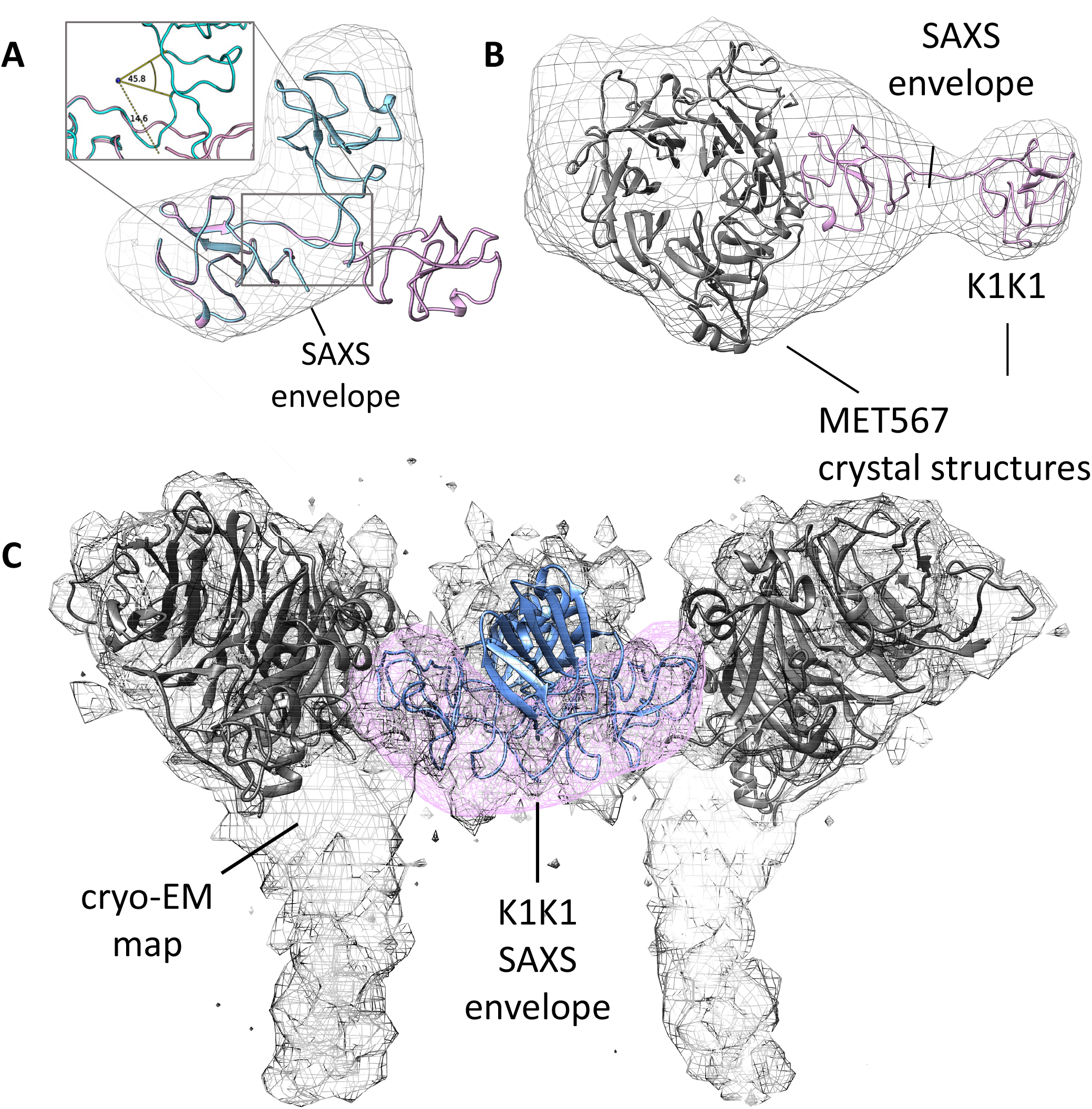
SAXS models for K1K1 alone and in complex with MET567 receptor fragment. (A) The measured SAXS envelope is not compatible with the elongated K1K1 crystal structure (pink) but perfectly accommodates a bend conformation of K1K1 (cyan). To fit the SAXS envelope, the linker region is bent by roughly 60° (B) The ab initio SAXS envelope of the 1:1 complex of K1K1 and MET567 receptor fragment shows a “pan-handle” extension, which accommodates K1K1. The CR domain is partially protruding from the bottom of the envelope and extra volume might be occupied by glycosylated side chains not modelled. Images generated with UCSF Chimera. (C) The SAXS envelope of K1K1 in solution (pink) overlaps well with part of the cryo-EM map (emd_23923.map) accommodating the two kringle domains of the NK1 dimer (blue). Cryo-EM map for 7mob.pdb.

### *In vitro* biological data

To analyse MET receptor activation and downstream signalling induced by K1K1, HeLa cells were stimulated with K1K1, K1K1H6, and monovalent K1H6 at different concentrations using human HGF/SF as a reference. MET receptor phosphorylation and activation of the key downstream signalling molecules Akt and ERK were confirmed by Western blot (Fig. 4A) in which both K1K1 and K1K1H6 stimulate phosphorylation of the pathway down to the lowest tested concentration of 100 pM. As predicted, treatment with K1H6 is indistinguishable from the negative control treatment, like what we observed with the chemically synthesised K1 domain previously (Ollivier *et al*, 2012). We then performed precise quantification of Akt and ERK activation in HeLa cells stimulated with HGF/SF, K1K1, K1K1H6, and K1H6 using AlphaScreen™ technology. For p-Akt, both K1K1H6 and K1K1 show similar phosphorylation in the low nanomolar range with no significant effect caused by the presence or absence of the poly-histidine tag. Compared to HGF/SF, both variants show lower maximum activation values of Akt (Fig. 4B). Interestingly, activation of the Ras-Raf-MAPK pathway as measured by the phosphorylation of ERK showed no significant differences between stimulation by HGF/SF, K1K1H6, or K1K1 at the tested concentrations. Again, no activity was observed with K1H6 even at the highest concentrations. We next confirmed the direct binding of K1K1H6 to recombinant human MET-Fc chimera, a soluble dimeric protein comprising the whole extracellular part of the MET receptor (MET922 fused to the Fc region of human IgG1) at different concentrations (Fig. 4D, S3A) using the AlphaScreen™ technology. To accurately determine the K_D_, we performed a full kinetic analysis of K1K1 binding to immobilized MET567 by surface plasmon resonance spectroscopy which furnished an apparent K_D_ of 205 nM (Fig. S3B), which is similar to the K_D_ obtained for NK1 (200 nM, Fig. S3C).

**Figure 4.**
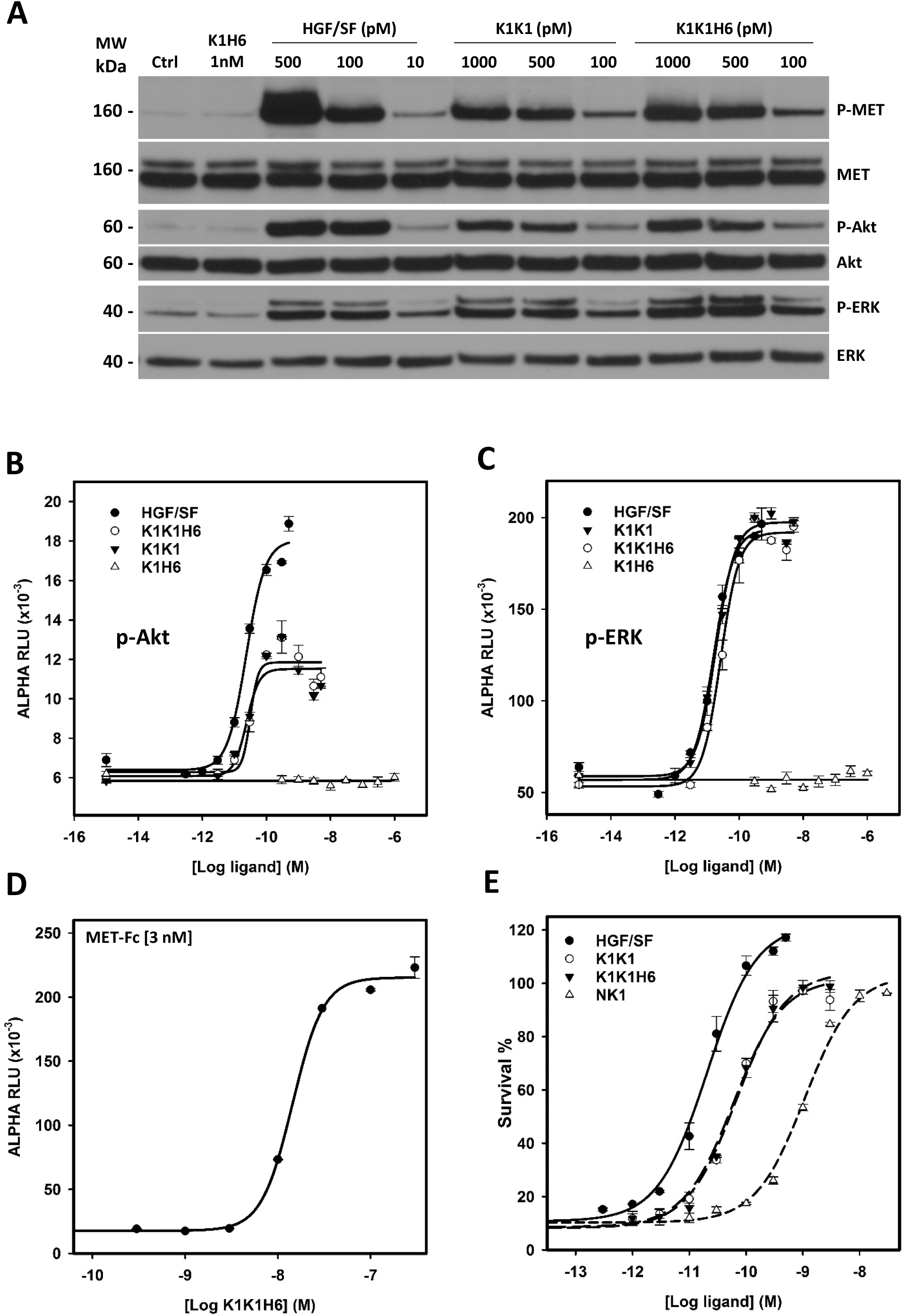
In vitro activity of the HGF/SF, K1K1, K1K1H6, and K1H6. (A) Phosphorylation analysis by Western blot on Hela cell lysates after stimulation with ligands for 10 min at concentrations indicated above each lane. Loading controls are based on total MET, total Akt, and total ERK present in each lane. (B) AlphaScreen™ measurements of p-Akt and (C) p-ERK activation in HeLa cells after 10 min stimulation with K1K1, K1K1H6, HGF/SF and K1H6. (D) Binding determination using the ALPHAScreen™ saturation binding assay. Seven concentrations of K1K1H6 were tested on several different concentrations of MET-Fc (AA 25-922) (Suppl. Fig S3A). Shown is the binding of K1K1H6 to 3 nM MET-Fc. (E) Ligand induced MDCK survival after overnight treatment with the apoptotic inducer anisomycin. Indicated is the percentage of viable cells compared to no-anisomycin treatment after exposure to HGF/SF, K1K1, K1K1H6, and NK1 at different concentrations.

Having established MET receptor binding and remarkably potent activation of the MET signalling pathway by both K1K1 and K1K1H6, we subsequently focused on demonstrating its potency in various phenotypic biological assays. We first performed a cell viability assay based on metabolic activity on MDCK cells incubated overnight (16 hours) in the presence of anisomycin, an apoptosis inducer, which prevents protein synthesis and leads to cell death. We looked at the cell survival after the addition of HGF/SF, K1K1, K1K1H6 and NK1 at different concentrations during anisomycin treatment. While HGF/SF was most effective, K1K1 and K1K1H6 were equals in preventing cell death and much more potent than native NK1 in this assay (Fig. 4E). We also used this assay to validate the specific activities of different preparations of K1K1 and K1K1H6 showing very little batch-to-batch variation and similar relative activity when compared to HGF/SF and NK1 (Fig. S4A) confirming the reliability of the protein production and purification protocol.

An MDCK scatter assay (Stoker & Perryman, 1985) was used as a sensitive phenotypical assay to determine the minimal concentration needed to activate MET and induce the epithelial-mesenchymal transition and cell scattering. K1K1 was able to exert an effect at a concentration down to 1 pM, ten times lower than native HGF/SF and one thousand times lower than NK1 (Fig. S4B).

To examine the role of the low-affinity heparin binding sites in the kringle domain, we used the analogues K1K1S2 and K1K1S4 with specific reverse-charge mutations altering heparin binding. These variants were compared with HGF/SF and K1K1 in an MDCK scattering assay (Fig. 5A). Both variants had reduced activity with observable scattering for K1K1S2 down to 300 pM and no scattering observed at 100 pM while K1K1 still showed activity at a concentration ten times lower (10 pM). K1K1S4 only showed activity in the high nanomolar range and was inactive at 1 nM. We further performed a 3D morphogenesis assay using MDCK cells (Fig. 5B and Fig. S5A and B) and confirmed the results obtained with the scatter assay, indicating that the reverse-charge mutations reduced the biological activity of K1K1S2 and K1K1S4. The loss of biological activity induced by the mutations was also observed in SKOV3 cells using a quantitative Boyden chamber migration assay (Fig. 5C). Finally, we examined the impact of the mutations on biological activity at the level of the MET receptor signalling pathway by investigating the phosphorylation status of MET, Akt, and ERK (Fig. S5C). Altogether these data indicate that K1K1S2 and K1K1S4 are weaker agonists than K1K1 by one and two orders of magnitude respectively and suggest that the low-affinity heparin binding site within the K1 domain plays a significant role in MET activation by K1K1.

**Figure 5.**
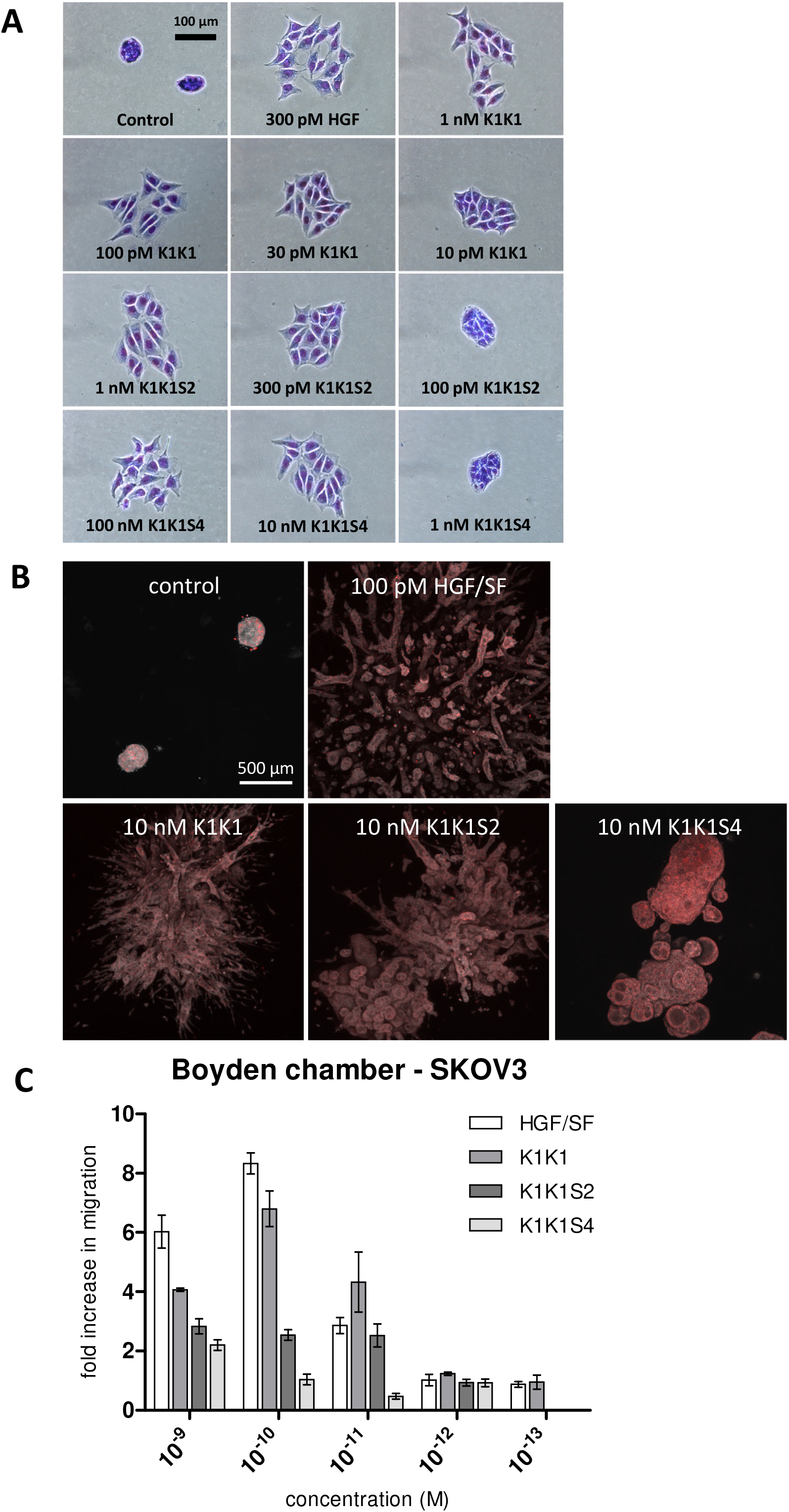
Effects on cell motility, migration, and morphology. (A) MDCK cell scattering at different concentrations K1K1, K1K1S2 and K1K1S4 showing the lowest concentration at which each protein is still active and the subsequent dilution at which no more scattering is observed. HGF/SF is used as positive control to generate maximum scattering. For complete half-log dilution of agonist series, see Data Source Image File. (B) 3D reconstruction by z-stacking of fluorescence microscopy images taken of large MDCK cell colonies stimulated with 100 pM HGF/SF or 10 nM K1K1 and mutants for four weeks. The combined fluorescence of DAPI (red) and Evans blue staining (grey scale) shows the extensive branching morphogenesis and tubulogenesis induced by both proteins. (C) Boyden chamber migration assay using SKOV3 cells. Cells were treated for 6 hours with indicated concentrations of HGF/SF, K1K1, K1K1S2 and K1K1S4. Migration is presented as fold increase over control. Error bars represent mean +/- SD based on quadruplicates (n=4).

### *In vivo* activation of MET signalling

Having established the superior potency of K1K1 over NK1 in different in vitro assays that sometimes matched or even surpassed native HGF/SF, we were keen to start several *in vivo* mouse studies to look at the effects of K1K1. We first validated that K1K1 is diffusible and biologically active *in vivo*, whatever the injection route (intraperitoneal (I.P.) versus intravenous (I.V.)), by measuring phosphorylation of the MET receptor, Akt, and ERK in liver homogenates 10 minutes after injection (Fig. S6). Since no significant difference was observed between I.P. and I.V. administration routes, all subsequent injections were done by I.P for experimental convenience. To study the effect of different doses, 8-week-old FVB mice were injected with different amounts of K1K1 (0.1 to 5 μg per 20 g animal) after which the animal was sacrificed 10 minutes later. The phosphorylation status of the MET signalling pathway was established in the liver homogenates. Control as well as 0.1 μg of K1K1 did not result in any detectable MET, Akt, and ERK phosphorylation signal compared to control while injection of 0.5, 1, and 5 μg gave uniform activation of the pathways (Fig. 6A). A second experiment was performed to determine the duration of the stimulation by sacrificing the animals at 10, 30, 60, and 90 minutes, using a K1K1 dose of 5 μg. The data obtained show a diminishing signal over time detectable up to 60 min (Fig. 6B) that could correspond to natural receptor desensitisation.

**Figure 6.**
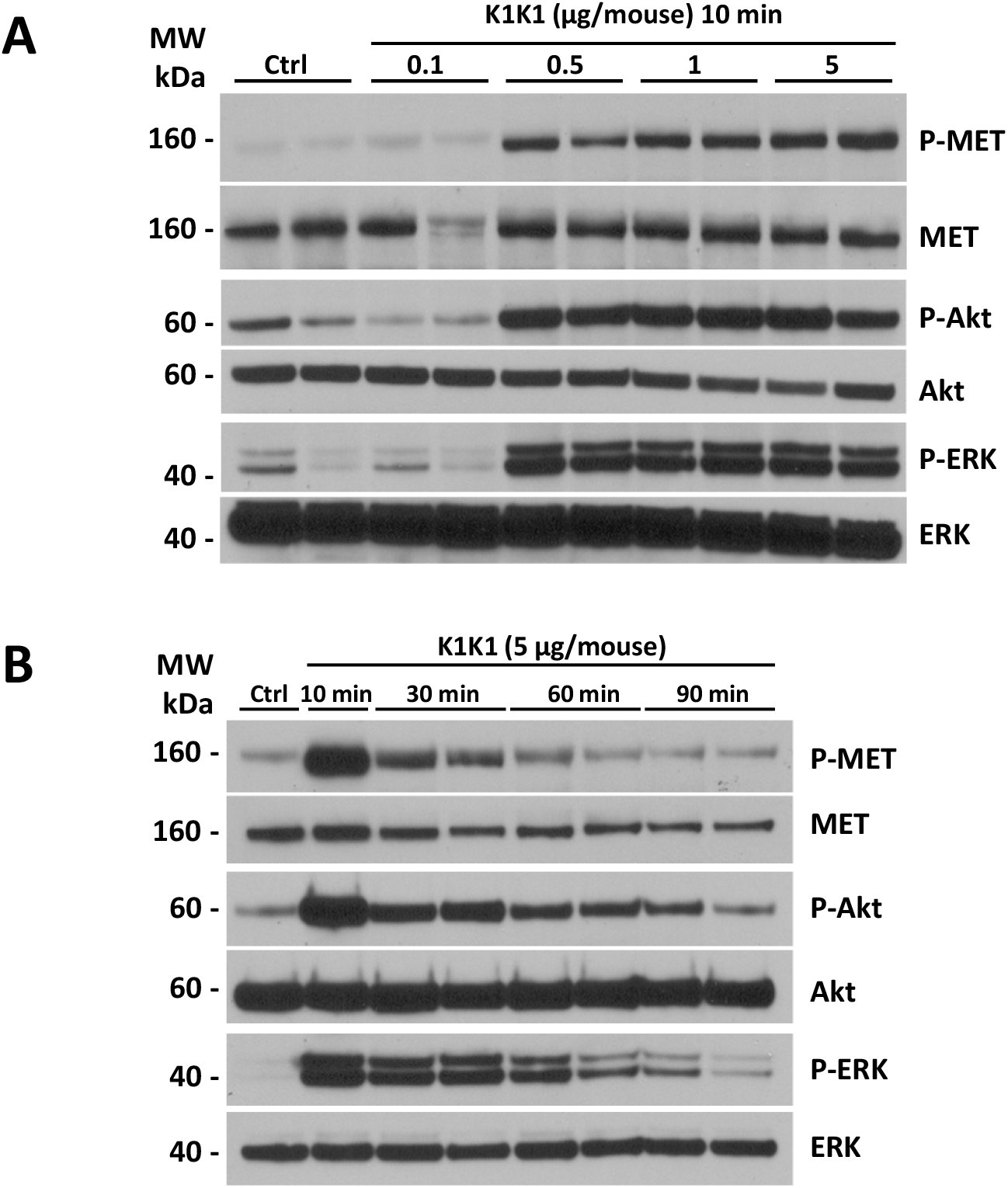
Dose response and MET pathway activation kinetics after intraperitoneal injection. (A) 8-week-old FVB mice were injected with PBS (Ctrl) or different amounts of K1K1 after which MET, AKT, and ERK phosphorylation in liver homogenate was determined using Western blot. Mice were sacrificed 10 minutes after injection. Results are presented as experimental duplicate (n=2). (B) MET, AKT, and ERK phosphorylation were detected by Western blot at different time points after injection of 5 μg of K1K1. Results are presented as experimental duplicate (n=2) except for control and 10 min conditions. Both blots present total MET, Akt and ERK proteins as loading controls.

### *In vivo* efficacy of K1K1 in the treatment of alcoholic steatohepatitis

To evaluate the *in vivo* efficacy of K1K1 applied to a liver disease model, we used a validated mouse model of subchronic alcohol exposure, the adapted Lieber DeCarli (LDC) model (Lieber & Decarli, 1989) in order to study steatosis, a common feature of several liver diseases.

As expected, alcohol consumption induced steatosis in liver of mice fed with alcohol (LDC + Vehicle group) which was not observed in mice fed with a control diet (Control + Vehicle) or a control diet with K1K1 treatment (Control + 10 μg K1K1) (Fig. 7A and Fig. S7A). The treatment with different doses of K1K1 (0.4, 2, and 10 μg) significantly decreased the steatosis in mice fed with alcohol (LDC + K1K1) (Fig. 7B). Regarding the factors involved in steatosis improvement, K1K1 treatment was able to significantly increase mRNA expression of ApoB, PPARα and LDLR (Fig. 7C). The highest doses of K1K1 (2 and 10 μg) also significantly decreased the mRNA expression of TNFα and IL-6 in LDC mice, while a moderate increase of both markers was observed in untreated mice compared to control health mice. Interestingly, while mRNA expression of MET decreased by ethanol exposure alone, it was induced by K1K1 treatment (Fig. S7B).

**Figure 7.**
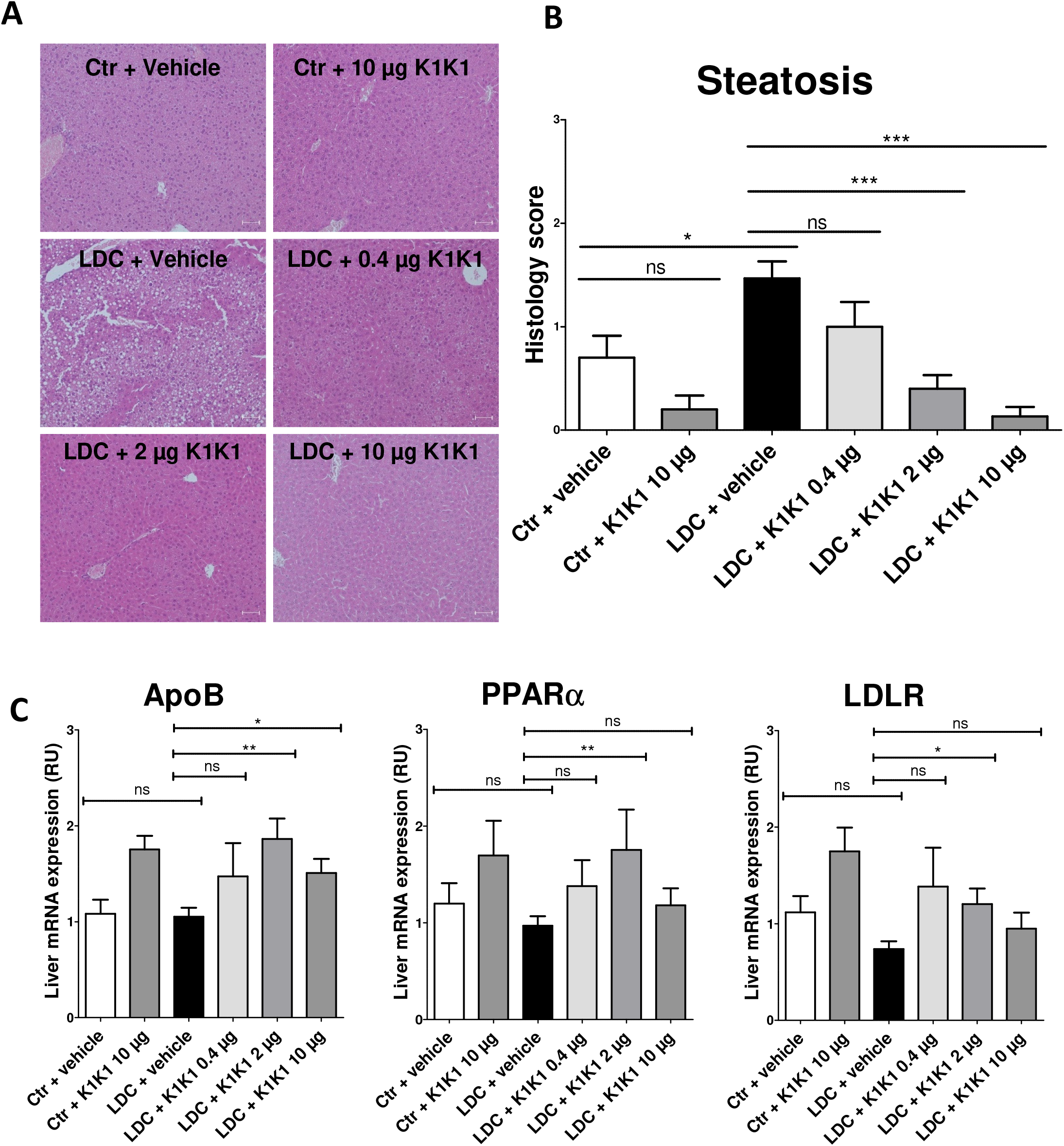
In vivo evaluation of the efficacy of K1K1 in a mouse model of alcoholic steatohepatitis. (A) Hematoxylin-erythrosin B staining of mouse livers submitted to an adapted Lieber DeCarli model (20x magnification). Analyses were performed on 10 animals in control groups (Ctr + Vehicle and Ctr +10 μg K1K1) and 15 animals in ethanol treated groups (LDC ± K1K1). Scale bar = 50 μm (B) Steatosis was quantified in each mouse liver using a steatosis score (see STAR Methods). Results are expressed as mean ± SD (C). The mRNA expression of triglyceride metabolism markers (ApoB, PPARa, and LDLR) was analysed in mouse livers using RT-qPCR with b-actin as housekeeping gene. Results are expressed in relative units (RU) and represented as mean ± SD.

Taken together these results show that K1K1 significantly improved steatosis in a mouse model of liver injury. Our data suggest that this therapeutic effect is mediated in part by the increase of protective factors such as ApoB and PPARα and the lowering of proinflammatory cytokines, induced by K1K1 treatment.

## Discussion

HGF/SF-MET signalling is essential for liver development (Schmidt *et al*, 1995; Uehara *et al*, 1995) and regeneration (Borowiak *et al*, 2004; Huh *et al*, 2004; Liu, 2004) and pre-clinical studies employing either recombinant HGF/SF (Horiguchi *et al*, 2009), HGF/SF gene delivery (Ueki *et al*, 1999) or HGF/SF-transfected mesenchymal stem cells (Moon et al., 2019) all demonstrated a strong therapeutic potential for HGF/SF in pre-clinical models of liver diseases. However, all failed to offer a robust platform toward the use of HGF/SF in the clinic. Indeed, both gene therapy and cell-based therapies meet considerable regulatory challenges and therapeutic applications of recombinant HGF/SF face strong and well-known limitations: the protein has a complex multidomain protein susceptible to proteolytic degradation, is unstable in physiological buffers, and exhibits limited tissue diffusibility and penetration. Hence, harnessing the HGF/SF-MET pathway in regenerative medicine will rely on engineered and improved HGF/SF derivatives in terms of stability and tissue availability.

Several molecules have been developed in recent years to overcome the limitations of HGF/SF in therapy. These included engineered variants of the natural growth factor (Cassano *et al*, 2008; Sinha Roy *et al*, 2010), agonistic monoclonal antibodies (Gallo *et al*, 2014; Kim *et al*, 2019), synthetic cyclic peptides (Sato *et al*, 2020), and aptamers (Ueki *et al*, 2020). All the engineered variants of HGF/SF developed thus far, however contain the high-affinity HSPG binding site present in the N-domain and thus share with HGF/SF the feature of limited diffusion and tissue penetration. The design of a new MET ligand with reduced affinity for HSPG, therefore was the prime objective of the present study. Building on available structures, mutagenesis data, and the biological activity of multimeric assemblies of the K1 protein (Simonneau *et al*, 2015), we rationalised that a K1K1 covalent dimer may have yielded a MET agonist with attractive physico-chemical and biological properties.

The structural data reported in this work show that K1K1 displays a perfect symmetrical positioning of the K1 units in its crystalized form and thus readily opens the possibility of a direct MET dimerization in a 1:2 K1K1:MET configuration. SAXS data indicate a bending of K1K1 when in solution, nicely fitting the natural position of the K1 domains within the NK1 dimer bound to MET as observed in recent cryo-EM structural analysis (Uchikawa *et al*, 2021). Thus, the structural data suggest that flexibility is important for enabling the proper positioning of K1 domains upon binding to MET.

K1K1 displayed a vastly superior stability in physiological buffers compared to native HGF/SF and a potency approaching activity at picomolar concentrations *in vitro* (Fig. 4A and Fig. 5A). Importantly, the improved diffusibility into tissues as a result of reduced binding to heparin (Fig S8) clearly is a further and major component of the remarkable activity displayed by K1K1 in the studies *in vivo* (Figs 6 and 7). SPR analysis confirmed a ten-fold reduction of heparin binding in K1K1 (K_D_ ~5.6 μM) compared to NK1 (K_D_ ~0.43 μM) while the affinity of the K1K1 mutants S2 and S4 was below the limits of sensitivity of SPR (Fig. S8). Interestingly, these mutants displayed reduced biological activity compared to K1K1 indicating that the presence of the low affinity site for heparan sulfate in the K1 domain plays a role in the potent agonistic activity *in vitro* as well as *in vivo*. Seen from a different angle, K1K1 can be regarded as a tool that can be easily manipulated and varied to investigate more in depth the role of heparan sulfate molecules in ligand-receptor binding and receptor activation, independently of the strong HS-binding N-domain.

Altogether, the structural, biochemical, and biological data reported here suggest that K1K1 might serve as a lead candidate for protein therapy of several human pathologies where activation of the HGF/SF-MET signalling axis can be beneficial. Activation of the MET pathway has been known to ameliorate alcoholic steatohepatitis as well as liver fibrosis or cirrhosis in pre-clinical models of liver disease (Matsuda, 1997; Matsuda *et al*, 1995; Tahara *et al*, 1999; Taïeb *et al*, 2002) and, conversely, deletion of the MET gene in postnatal mice causes hyperlipidemia, severe steatosis, and progressive liver disease (Kroy *et al*, 2014). No specific drug has been approved as yet for the treatment of alcoholic and non-alcoholic liver disease (Sharma *et al*, 2020) but we submit that the K1K1 protein data presented here may emerge as a strong candidate for clinical development owing to its favourable physical chemistry and its strong activity in a well-established clinical mouse model of alcoholic steatohepatitis (Fig. 7 and S7A).

Finally, we wish to emphasize that HGF/SF is a broad-acting morphogen that controls development and regeneration of other major epithelial organs beyond liver: HGF/SF - MET signalling is essential for lung alveologenesis (Calvi *et al*, 2013) and for kidney development (Ishibe *et al*, 2009). Thus, we surmise that potent MET agonists may find many applications beyond liver disease notably in the large number of patients with acute and chronic kidney damage (Zhou *et al*, 2013) and chronic lung emphysema (Shigemura *et al*, 2005) and may well find an urgent application in reducing lung damage and fibrosis in patients suffering from COPD (chronic obstructive pulmonary disease) or severe COVID-19 (George *et al*, 2020; Zuo *et al*, 2010).

## Materials and Methods

### Cloning of K1K1 and K1K1 variants

The cDNA encoding K1K1 and K1K1H6 was generated using a fusion PCR reaction after separate PCR amplification of two K1 domains from a wild-type human NK1 template. The forward primer, ATCATCCCAT GGCCATTAGA AACTGCATCA TTGGTAAAGG ACG, was used to amplify the first kringle domain introducing a 5’ NcoI restriction site. The reverse primer for the first K1, TTCAACTTCT GAACACTGAG GA, was used to introduce the linker sequence S E V E placed in-between the two K1 domains. S E V E is naturally found in HGF/SF in between the K1 and K2 domains. The forward primer for the second K1 domain, TCAGAAGTTG AATGCATCAT TGGTAAAGGA CG, introduces the fusion-overlap with the SEVE linker sequence. A reverse primer with or without a six-histidine tag, ACAGCGGCCG CTCATCAATG ATGATGATGA TGATGTTCAA CTTCTGAACA CTGAGGA and ACAGCGGCCG CTCATCATTC AACTTCACTA CACTGAGGAA T respectively, introduced two stop codons followed by a NotI restriction site. Following the fusion PCR reaction, the product was digested with NcoI and NotI and ligated into the pET45b(+) expression plasmid (Novagen/EMD Millipore). Integration in the pET45b(+) MCS introduced the additional amino acid sequence M A I R N upstream of the first cysteine of K1 domain. The heparin mutants S2 and S4 were based on existing kringle-variant NK1 templates. These templates introduced the reverse-charge amino acid substitutions K10E, R12E, K93E, R95E for K1K1S2 and K10E, R12E, K48E, R59E, K93E, R95E, K131E, R142E for K1K1S4.

### Protein expression

Recombinant human NK1 protein (residues 28-209) was expressed in Pichia pastoris and purified as described in Chirgadze et al., 1999 (Chirgadze et al., 1999). Recombinant human MET567 was expressed in CHO Lec 3.2.8.1 cells and purified as described in Gherardi et al., 2003 (E Gherardi et al., 2003). Human HGF/SF was stably expressed in NS0 myeloma cells and purified as described in Gherardi et al., 2006 (Ermanno Gherardi et al., 2006).

For the expression of K1K1 and K1K1 variants, a single colony of freshly transformed E. coli BL21(DE3) was used to inoculate an overnight 5 mL Luria broth (LB) culture containing 100 μg/mL of ampicillin. This culture was used to inoculate 500 mL of LB with ampicillin shaking at 250 rpm and grown to an optical density (600 nm) of 0.6-0.8 at 37 °C. The culture was then cooled down to 18 °C while shaking after which IPTG was added to a final concentration of 0.1 mM. The culture was grown at 18 °C with shaking for 24 h after which the bacterial cells were harvested by centrifugation for 30 min at 10,000 rcf at 4 °C and the cell pellet was stored at −80 °C.

### K1K1 extraction from inclusion bodies

The frozen bacterial cell pellet was thawed on ice and resuspended in 25 mL 50 mM Tris pH 8.5, 500 mM NaCl, with the addition of one tablet of cOmplete™ EDTA-free Protease Inhibitor Cocktail (Roche), 1 unit of Pierce Universal Nuclease and 10 μg of lysozyme. The suspension was incubated rotating at 4 °C for 30 min before being placed on ice and sonicated using ten pulses of 30 s with 60 s pause in between using a Sonic Ruptor 400 (Omni International) with a OR-T-750 tip at 100% power output. The suspension was centrifuged at 10,000 rcf for 10 min, the supernatant discarded, and the pellet resuspended in 25 mL 50 mM Tris pH 8.5, 500 mM NaCl with 0.4% Triton X-100 using a glass Potter-Elvehjem tube and PTFE pestle. The suspension was incubated at 4 °C rotating for 30 min after which it was centrifuged at 10,000 rcf for 10 min, the supernatant discarded, and the pellet resuspended in 25 mL 50 mM Tris pH 8.5, 500 mM NaCl with 0.025% NP40 using the Potter-Elvehjem tube and pestle. The suspension was again incubated at 4 °C rotating for 30 min, centrifuged for 10 min at 10,000 rcf, the supernatant discarded, and the pellet resuspended in 25 mL 50 mM Tris pH 8.5, 500 mM NaCl using the Potter-Elvehjem tube and pestle. The suspension was once more incubated at 4 °C rotating for 30 min, centrifuged at 10,000 rcf for 10 min, and the supernatant discarded. The final pellet was resuspended in 20 mL 50 mM Tris pH 8.5, 500 mM NaCl, 2 M arginine, 0.5 mM GSSG, 5 mM GSH. This suspension was incubated for three days at 4 °C on a rotary wheel.

### K1K1 purification

After three days of incubation in Tris buffer with arginine, the inclusion body suspension was centrifuged at 20,000 rcf for 30 min at 4 °C. The supernatant was transferred to a new centrifuge tube and centrifuged again at 20,000 rcf for an additional 30 min at 4 °C. Unless a pellet was visible, in which case a third centrifugation was performed in a new tube, the cleared supernatant was diluted 100 times in 2 L of 50 mM Tris pH 7.4, 150 mM NaCl and filtered. The diluted supernatant was loaded onto a 5 mL Heparin HiTrap™ column (GE Healthcare) and eluted with a gradient up to 1 M NaCl, in 50 mM Tris pH 7.4. Peak fractions were pooled, concentrated, and loaded on a HiLoad^®^ 16/600 Superdex 200 pg column (GE Healthcare) equilibrated in 50 mM Tris pH 7.4, 500 mM NaCl. Peak fractions were pooled and concentrated to 5 mg/mL before being used or stored after flash freezing. For K1K1S2 and K1K1S4, instead of Heparin HiTrap™ column, the first step purification was done on a 5 mL HisTrap™ HP column (GE Healthcare).

### Protein crystallization and x-ray diffraction

For protein crystallization, K1K1 and K1K1H6 were dialyzed in 10 mM Tris pH 8.5, 100 mM NaCl and concentrated to 12 mg/mL and 11.8 mg/mL respectively. A pre-crystallization screen (PCT, Hampton Research) was performed to confirm these concentrations were favourable for crystallization after which 48-well sitting-drop plates (Hampton Research) were set up using Crystal screen 1 (Hampton Research), JCSG-plusTM (Molecular Dimension) and Morpheus^®^ (Molecular Dimensions) crystallization screens. After overnight incubation at 17 °C, the first crystals of K1K1H6 appeared in the Morpheus^®^ screen in condition C7 consisting of 100 mM MOPS/HEPES pH 7.5, 30 mM sodium nitrate, 30 mM sodium phosphate dibasic, 30 mM ammonium sulphate, 20% v/v glycerol, 10% w/v PEG4000. Further optimization was performed with different protein to crystallization solution ratios (1:2, 1:1, 2:1) in a 24-well plate using hanging drop. After 24 h, good crystals were obtained in a 2:1 protein-to-solution ratio condition. In addition to the cryo-protection intrinsic to the Morpheus□screen, additional glycerol was added up to 25% of the final volume before crystals were collected using a CryoLoop (Hampton Research) and flash frozen and stored in liquid nitrogen. Crystallizations of K1K1 went through a similar optimization procedure and yielded good crystals in a condition of 100 mM Tris/Bicine pH 8.5, 30 mM Sodium nitrate, 30 mM Sodium phosphate dibasic, 30 mM Ammonium sulfate, 12.5% w/v PEG 1000, 12.5% w/v PEG 3350, 12.5% v/v MPD. Data was collected at the ESRF in Grenoble, France, at beamline at BM14 for K1K1H6 and ID23-1 for K1K1 and was solved by molecular replacement using the NK1 kringle domain taken from 1BHT.pdb (Ultsch et al., 1998) as a search model resulting in a structure with a resolution of 1.7Å and an Rwork/Rfree of 19.7/22.7% for K1K1 and a structure with a resolution of 1.8Å and an Rwork/Rfree of 16.5/19.4% for K1K1H6. Both proteins crystalized in monoclinic space group P1 21 1.

### Crystal structure determination

For both K1K1H6 and K1K1, the MTZ file generated by the ESRF EDNA framework Fast Processing System was used, downloaded through the ISPyB server. At the different sessions of measurement, the EDNA auto processing used AIMLESS (v0.3.3) and POINTLESS (v1.9.8) to create the MTZ file for K1K1 while for K1K1H6 the MTZ file was generated using AIMLESS (v0.5.2) and POINTLESS (v1.9.25). Molecular replacement was performed using Phaser (v2.8.3) the Phenix software package (v1.19.2) (Liebschner *et al*, 2019) using the kringle domain from 1BHT.pdb (Ultsch et al., 1998) followed by several rounds of refinement using Phenix refine (Afonine et al., 2012) and WinCoot (Emsley *et al*, 2010) (v0.9.4.1). The dataset of K1K1 and K1K1H6 had an upper resolution limit of 1.61 and 1.39 respectively, a resolution cut-off was set to 1.8 for K1K1 and 1.7 for K1K1H6. The data collection and refinement parameters and statistics are given in table 1. The data collection and refinement statistics are given in Supplementary table S1. The molecular graphics presented in this manuscript were generated with PyMOL (v2.4.1) and Chimera (v1.15) (Pettersen *et al*, 2004).

### Small angle X-ray scattering (SAXS)

Solution scattering data were collected at ESRF BM29 using a sec-1 frame rate on Pilatus 1 M detector located at a fixed distance of 2.87 m from the sample, allowing a global q range of 0.03-4.5 nm with a wavelength of 0.01 nm. SEC-SAXS experiments were carried out using Nexera High Pressure Liquid/Chromatography (HPLC; Shimadzu) system connected online to SAXS sample capillary. For these experiments, 35 μL of each sample at the concentrations indicated in the Table were injected into a Superdex 200 PC 3.2/300 Increase column (GE Healthcare), pre-equilibrated with 25 mM Tris pH 7.4, 150 mM NaCl. Frames corresponding to protein peaks were identified, blank subtracted and averaged using CHROMIXS2. Radii of gyration (Rg), molar mass estimates and distance distribution functions P(r) were computed using PRIMUS in the ATSAS package (v3) (Petoukhov *et al*, 2012). Modelling of flexible loops and glycosylation were carried out using COOT, CORAL and SASREF. Comparison of experimental SAXS data and 3D models from structural models was performed using CRYSOL. A summary of SAXS data collection and analysis results is shown in Supplementary Table S2.

### AlphaScreen™ MET binding assay

Saturation assays for binding of K1K1H6 to recombinant MET-Fc protein were performed in 384-well microtiter plates (OptiPlate™-384, PerkinElmer©, CA, USA, 50 μL of final reaction volume). Final concentrations were 0-300 nM for K1K1H6, 0-10 nM for hMET-Fc (8614-MT-100, R&D Systems), 10 μg/mL for Ni-NTA coated donor beads and protein A-conjugated acceptor beads. The buffer used for preparing all protein solutions and the beads suspensions was PBS (10 mM phosphate buffer pH 7.4, 148 mM NaCl, 2 mM KCl), 5 mM HEPES pH 7.4, 0.1% BSA. K1K1H6 (10 μL, 0-1.5 μM) was mixed with solutions of MET-Fc (10 μL, 0-50 nM). The mixture was incubated for 10 min (final volume 15 μL). Protein A-conjugated acceptor beads (#6760617C, PerkinElmer©; 10 μL, 50 μg/mL) were then added to the vials. The plate was incubated at 23 °C for 30 min in a dark box. Finally, Ni-NTA coated donor beads (6760619C, PerkinElmer©; 10 μL, 50 μg/mL) were added and the plate was further incubated at 23 °C for 60 min in a dark box. The emitted signal intensity was measured using standard Alpha settings on an EnSpire^®^ Multimode Plate Reader (PerkinElmer). Measurements are expressed as technical duplicates (mean+/- SD, n=2). The data corresponding to the 3 nM MET condition was subjected to a non-linear regression analysis using four-parameter sigmoidal equation using SigmaPlot software (v13 and v14.5).

### Cell culture

Madin-Darby Canine Kidney (MDCK, kind gift of Dr Jacqueline Jouanneau, Institut Curie, Paris, France) and Human cervical cancer HeLa cells, purchased from ATCC^®^ (American Type Culture Collection, Rockville, MD, USA), were cultured in DMEM medium (Dulbecco’s Modified Eagle’s Medium, Gibco, Karlsruhe, Germany), supplemented with 10% FBS (Fetal Bovine Serum, Gibco^®^, Life technologies, Grand Island, NY, USA) and 1/100 of ZellShieldTM (Minerva Biolabs GmbH, Germany). 24 h before treatment, the medium was exchanged with DMEM containing 0.1% FBS, and cells were then treated for subsequent experiments. SKOV-3 cells were cultured in RPMI 1640 medium supplemented with 10% FBS and penicillin-streptomycin all purchased from Gibco/ThermoFisher Scientific.

### Western Blots

The assay was performed according to Simonneau et al., 2015 (Simonneau et al., 2015). Hela cells were collected by scraping and lysed on ice with a lysis buffer (20 mM HEPES pH 7.4, 142 mM KCl, 5 mM MgCl_2_, 1 mM EDTA, 5% glycerol, 1% NP40 and 0.1% SDS) supplemented with freshly added protease inhibitor (1/200 dilution, #P8340, Sigma Aldrich) and phosphatase inhibitor (1/400 dilution, #P5726, Sigma Aldrich). Lysates were clarified by centrifugation (20,000 × g, 15 min) and protein concentration was determined (BCA protein assay Kit, Pierce^®^, Thermo scientific, IL, USA). The same protein amount (usually 20-30 μg) of cell extracts was separated by NuPAGE gel electrophoresis (4-12% Midi 1.0 mm Bis-Tris precast gels, Life technologies) and electrotransferred to polyvinylidene difluoride (PVDF) membranes (Merck Millipore). Membrane was cut between 80 and 110 kDa marker and at 50 kDa to probe simultaneously phospho or total MET, Akt and ERK. Membranes were probed overnight at 4 °C with primary antibodies diluted to 1/2000 in 5% BSA and 0.1% sodium azide in PBS using specific total MET (#37-0100 Invitrogen), total ERK2 (#SC-154 Tebu-bio), phospho-MET (Y1234/1235, clone CD26, #3077 Cell Signaling), phospho-Akt (S473, clone CD9E, #4060 Cell Signaling), phospho-ERK (T202/Y204, clone E10, #9106 Cell Signaling). After extensive washing with PBS-0.05% Tween^®^ 20 followed incubation with anti-mouse (#115-035-146) or anti-rabbit (#711-035-152) peroxidase-conjugated IgG secondary antibodies (Jackson ImmunoResearch) diluted to 1/30000 in PBS-casein 0.2%. Protein-antibody complexes were visualized by chemiluminescence with the SuperSignal^®^ West Dura Extended Duration Substrate (Thermo scientific) using a LAS-4000 imaging system (GE HeathCare Life Sciences) or X-ray films (CL-XposureTM Film, Thermo scientific).

### AlphaScreen™ phosphorylation assay

Measurement of ERK1/2 (T202/Y204) and Akt1/2/3 (S473) activation were performed on HeLa cells using AlphaScreen™ SureFire Kits (#TGRES500 and #TGRA4S500) according to manufacturer protocol with minor modifications. Into a 96 well plate (#353072, Flacon, Corning), 10 000 HeLa were seeded in 200 μL of DMEM supplemented with 10% FBS and let attached and grown for 24 h. Cells were next starved in DMEM supplemented with 0.1% FBS for 2 h. Cells were treated for 8 min with MET agonist from 0.3 to 500 pM with HGF/SF, from 3 pM to 50 nM with K1K1 or K1K1H6 and from 3 to 1000 nM with K1H6. Cells were quickly washed in cold PBS and AlphaScreen™ lysis buffer was added for 15 min under middle shaking (600 rpm on orbital rocker, MixMate, Eppendorf). Immediately, 5 μL of lysate in analyzed by addition of acceptor and donor bead mixes for 2 h at 23 °C. The emitted signal intensity was measured using standard Alpha settings on an EnSpire^®^ Multimode Plate Reader. Measured are expressed has experimental duplicates (mean+/- SD, n=2). The data were subjected to a non-linear regression analysis using four-parameter sigmoidal equation using SigmaPlot^®^ software (v13).

### Surface plasmon resonance (SPR) analysis

Affinity constants were measured using a Biacore T200 (GE Healthcare) at 25 °C using PBST (PBS + 0.05% Tween 20) as running buffer and a flow rate of 30 μL/min. For the measuring of affinity for MET, the receptor fragment MET567 comprising the N-terminal SEMA domain and the cysteine-rich/PSI domain was immobilized at low-density (515 RU) using amine-coupling on a CM5 sensor chip (GE Healthcare). A blank inactivated reference channel was used to correct for non-specific binding. 300 seconds injections were done using serial-diluted samples of K1K1, NK1, K1K1S2, and K1K1S4 testing a high range of 2 μM-0.125 μM and low range of 0.8 μM-12.5 nM injecting for 300 seconds to reach equilibrium. For the measurement of affinity for heparin, a heparin-coated SC HEP0320.a chip (Xantec) was used. With no reference channel available, all four channels were used for analysis. K1K1, NK1, K1K1S2, and K1K1S4 were injected in a wide range of concentrations (10 μM to 156 nM) for 300 seconds. Biacore Evaluation software version 3.2 was used for calculating affinity constants. Curves were plotted using Graphpad Prism (v5.04).

### SKOV-3 cell migration assays

Agonist-induced cell migration was measured by seeding 50,000 SKOV-3 cells resuspended in serum-free RPMI supplemented with 0.1% BSA (BioFroxx) in each of the upper wells of a 96-well Boyden chamber (AC96, Neuro Probe). HGF/SF, K1K1, K1K1S2, and K1K1S4 was loaded in the lower compartment and cells were left to migrate in a humidified 37 °C incubator with 5% CO_2_ for 6 h. Afterwards, non-migrated cells were removed, the cells migrated over the collagen coated (100 μg/mL Purecol, Nutacon) 8 μm pore-size membrane (Neuro Probe) were fixed in 4% paraformaldehyde in PBS and stained using HCS CellMask™ Green stain (ThermoFisher Scientific). Fluorescent intensity was measured using an Azure C600 (Azure Biosystems) and migration was quantified after control-adjustment and charted in Graphpad Prism.

### Madin-Darby canine kidney (MDCK) scatter assay

The assay was performed according to Stoker et al. (Stoker *et al*, 1987). MDCK cells were seeded at low density (2,000 cells/well on a 12-well plate, #353043 Falcon, Corning) in DMEM supplemented with 10% FCS to form compact colonies. After treatment, when colony dispersion was observed and the cells were fixed and colored by Hemacolor^®^ stain (Merck, Darmstadt, Germany) according to the manufacturer’s instructions. Representative images were captured using a phase contrast microscope with 40× and 100× magnification (Nikon Eclipse TS100, Tokyo, Japan).

### Morphogenesis Assay

Wells of a black thin clear bottom 24-well plate (#4ti-0241, Brooks Lifesciences) were coated with 200 μL of 2 mg/mL type 1 rat tail collagen (354249, Corning/BD Biosciences) diluted in DMEM equilibrated in 1/10 v/v of 7.5% w/v sodium bicarbonate solution (#25080-060, Gibco, ThermoFisher). Cells (500-700) were seeded into a 500 μL layer of a 1:1 collagen/Growth Factor Reduced MatrigelTM (#354230, Corning/BD Biosciences) mixed in DMEM balance with bicarbonate and were treated twice a week for one month. The cells were then fixed for 24 h with 4% paraformaldehyde in PBS, coloured for 3 h with 0.1 mg/L Evans Blue in PBS (#E2129, Sigma Aldrich), extensively washed in PBS and finally nuclei stained with 3 nM DAPI is PBS. Cell were observed using inverted microscope observed and confocal microscope (LSM880 Zeiss) with long distance 10× objective using 488 nm excitation Argon laser and LP 500-750 nm emission filter. 3D reconstitutions (14 μm slices) and maximum intensity projections of z-stack were realized with Zen software (v8.1).

### Survival Assay

In a clear bottom 96-well plate (#353072, Flacon, Corning), 12,500 MDCK cells were seeded and incubated for 24 h in DMEM supplemented with 10% FBS. The culture media was next added with increasing concentrations of the HGF/SF, K1K1, K1K1H6 and NK1 agonists together with the apoptosis inducer anisomycin (1.4 μM) for 15 h. The cells were then washed with PBS to eliminate dead cells and then incubated for 1 h in 200 μL of HBSS (#14025092, Gibco, ThermoFisher) containing 100 μM of Resazurin (#B21187, Sigma Aldrich). Fluorescence was then measured with Envision multimode reader (Ex: 560 nm, Em: 590 nm) with maximal signal gain set (90%) on control cells wells and on top/top reading configuration. The data were normalized to percentage of control cells signal and subjected to a non-linear regression analysis using four-parameter sigmoidal equation with minimum=0 and maximum=100 using SigmaPlot software (v13 and v14.5).

### *In vivo* MET activation in liver

All experimental procedures were conducted with the approval of the Ethics Committee for Animal Experimentation of the Nord Pas de Calais Region (CEEA 75). For kinetic analysis, 8-week-old FVB mice weighing 19-21 g (Charles River) were intraperitoneally injected with K1K1 (5 μg/mouse) and sacrificed after 10, 30, 60 or 90 min of treatment. For dose-response analysis, 8-week-old FVB mice weighing 19-21 g (Charles River) were intraperitoneally injected with 0.1, 0.5, 1 and 5 μg of K1K1 diluted in 100 μL of PBS and sacrificed per cervical dislocation 10 min after injection. To analyze the effect of the route of administration, 12-week-old C57BL/6 NRJ mice weighing 19-21 g (Janvier Labs) were intraperitoneally or intravenously injected with K1K1 (5 μg/mouse) and sacrificed after 10 min of treatment. Livers were immediately perfused with PBS supplemented with protease and phosphatase inhibitors, collected and then rapidly frozen in liquid nitrogen for subsequent protein extraction and Western blot analysis.

### Liver steatosis model

Eighty C57BL/6J female mice of 8 to 10 weeks (~ 19 g) provided by Janvier Labs were used in this study. They were housed in temperature- and humidity-controlled rooms, kept on a 12 h light-dark cycle. Animal procedures were conducted in accordance with French government policies. Mice were fed with a control liquid diet of adapted Lieber DeCarli ad libitum during a 7-day habituation process. Body weight and liquid diet consumption were monitored every two days. Mice were then separated in 6 groups:

– LDC Control Diet-fed mice + vehicle (n=10, Vehicle: 0.0 μg)
– LDC Control Diet-fed mice + product (n=10, highest dose: 10 μg)
– LDC OH-fed mice + vehicle (n=15, Vehicle: 0.0 μg)
– LDC OH-fed mice + product (n=15, Dose 1: 0.4 μg)
– LDC OH-fed mice + product (n=15, Dose 2: 2.0 μg)
– LDC OH-fed mice + product (n=15, Dose 3: 10 μg)

During a 10-days alcoholization period, mice received 5 intraperitoneal injections of 100 μL of sterile PBS or different doses of K1K1 diluted in sterile PBS. Ethanol-fed groups had unlimited access to adapted LDC containing ethanol and control mice were pair-fed with isocaloric control diet over this feeding period. On the 11th day, mice were sacrificed.

### Hematoxylin-erythrosine B staining

The livers embedded in paraffin and sectioned at 4 μm thick were deparaffinized and rehydrated in successive baths of xylene, absolute ethanol, ethanol 95% and tap water. The slides were stained with hematoxylin (Harris), rinsed, and stained with erythrosine B 0.8% (Sigma). The slides were finally dehydrated and mounted with Eukitt (GmbH). Slides were analyzed on a Zeiss Axio Imager M2 microscope. For histological evaluation, a score between 0 and 3 (3 = 75% steatosis) for steatosis was assigned to each section of the liver (3 areas per section).

### Real-time quantitative RT-PCR

Livers in 350 μL of RA1 and 3.5 μL of β-mercaptoethanol were mixed and frozen at −80 °C until RNA extraction. RNA extraction was carried out using the nucleospin RNA kit (Macherey-Nagel). The RNAs were quantified using a Nanodrop and the retro-transcription made using the high-capacity cDNA reverse transcription kit (Thermo Fisher). qPCRs were done by mixing 3.25 μL of H2O RNAse free from the RNA extraction kit to 6.25 μL of Fast SYBR green Master mix (ThermoFisher) and 0.25 μL of reverse and 0.25 μL of forward primers (10 pM) diluted 10x in distilled water. qPCRs were performed in a StepOne plus thermocycler (Thermo Fischer). The sequences of the primers used in the study are:

β-actin F: CCATGTCGTCCCAGTTGGTAA
β-actin R: GAATGGGTCAGAAGGACTCCTATGT
IL-6 F: GTTCTCTGGGAAATCGTGGA
IL-6 R: CAGAATTGCCATTGCACAAC
TNFα F: TGGGAGTAGACAAGGTACAACCC
TNFα R: CATCTTCTCAAAATTCGAGTGACAA
MET F: GACAAGACCACCGAGGATG
MET R: GGAGCCTTCATTGTGAGGAG
ApoB F: TACTTCCACCCACAGTCCC
ApoB R: GGAGCCTAGCAATCTGGAAG
LDLR F: TTCAGTGCCAATCGACTCAC
LDLR R: TCACACCAGTTCACCCCTCT
PPARα F: AGGCAGATGACCTGGAAAGTC
PPARα R: ATGCGTGAACTCCGTAGTGG

Results were presented as levels of expression relative to that of controls after normalizing with β-actin mRNA using the comparative Ct method.

## Supporting information

image source data

## Acknowledgments

We thank SATT-Nord (France) for financing the proof-of-concept experiments, Thierry Chassat from the PLETHA animal Facility in Lille Pasteur Institute for helpful advises and kind availability. Work in the laboratory in Pavia was made possible through funding from the Italian Ministry for Universities and Research (Departments of Excellence initiative) and from the Leukemia Research Fund (AIL Trentino). We would like to acknowledge Dr Maristella Magi (University of Pavia) for help at the ESRF beamline, Dr Claudia Scotti (University of Pavia) and Dr Dimitri Y. Chirgadze (University of Cambridge, UK) for help with structural analysis of K1K1 and K1K1H6, Maria Cristina Barbieri for technical support and Dr Raymond Pierce for proofreading.

