## Supplemental figures for "Dimerization of kringle 1 domain from hepatocyte growth factor/scatter factor provides a potent minimal MET receptor agonist"

### Supplementary figure S1

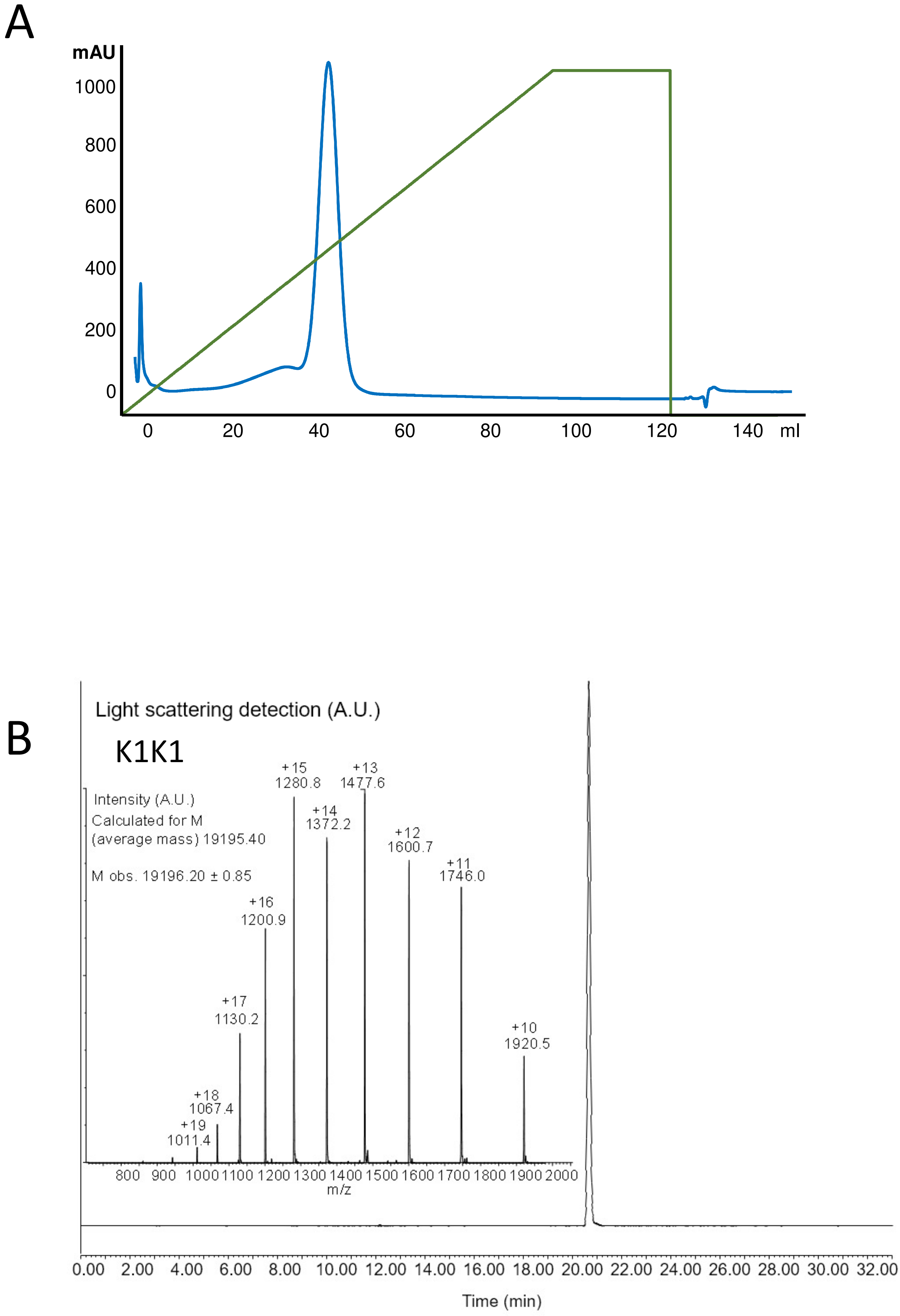

#### Supplementary figure S1

C

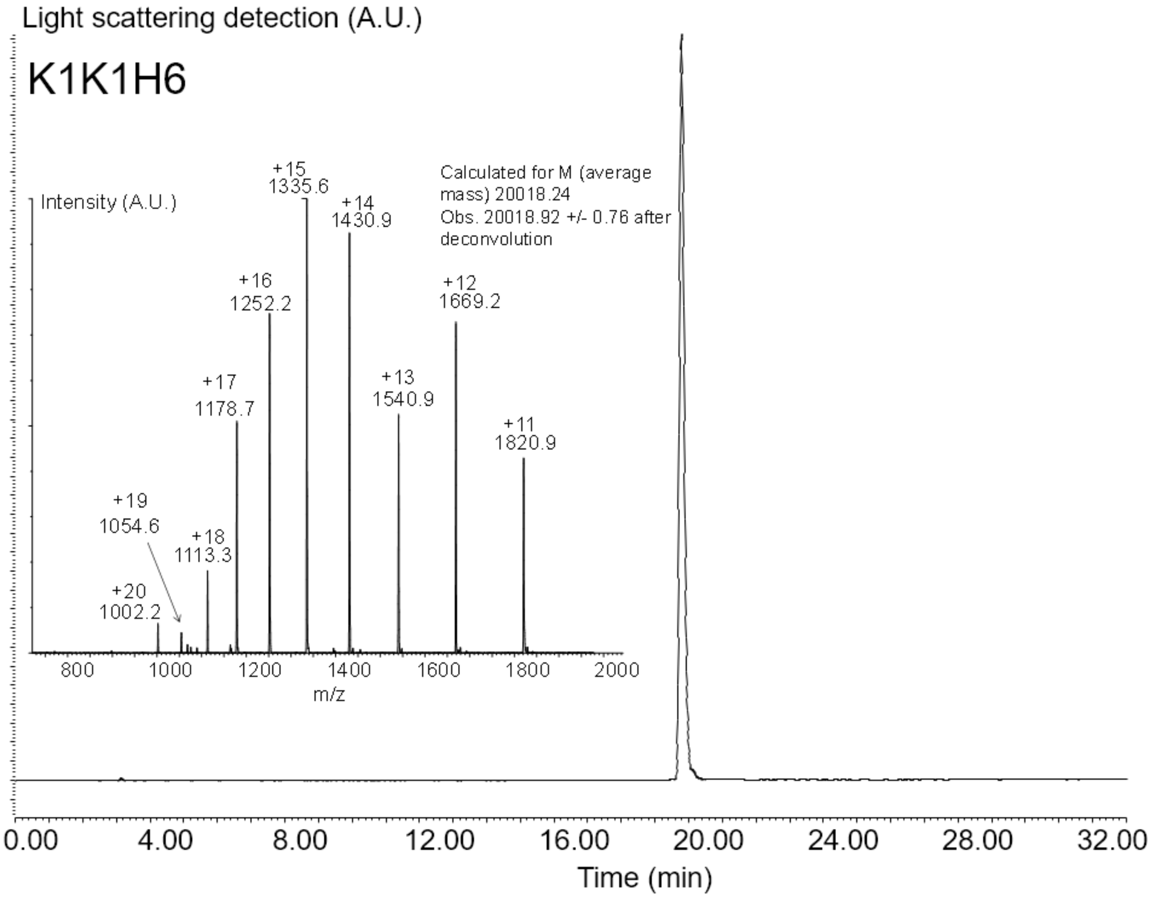


D

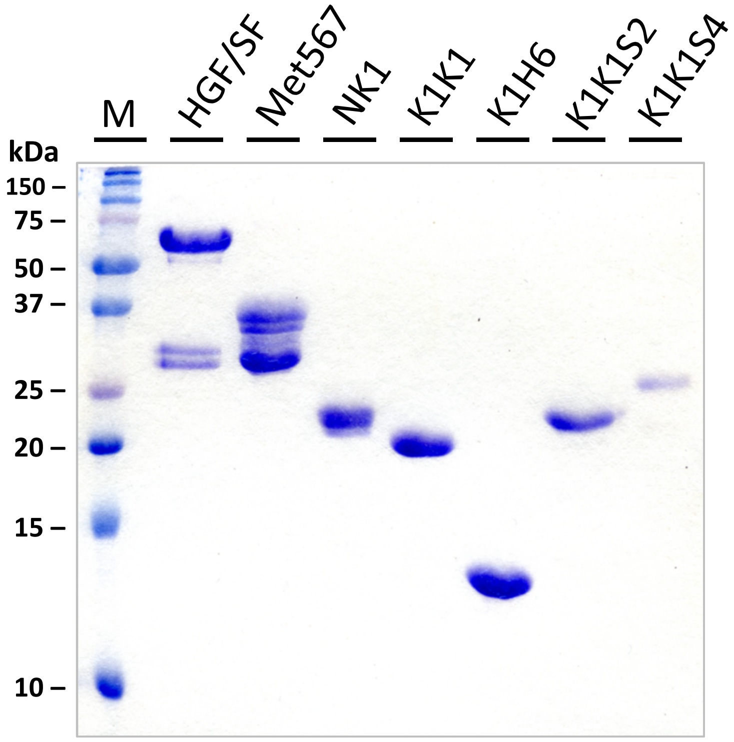

**Supplementary figure S2**

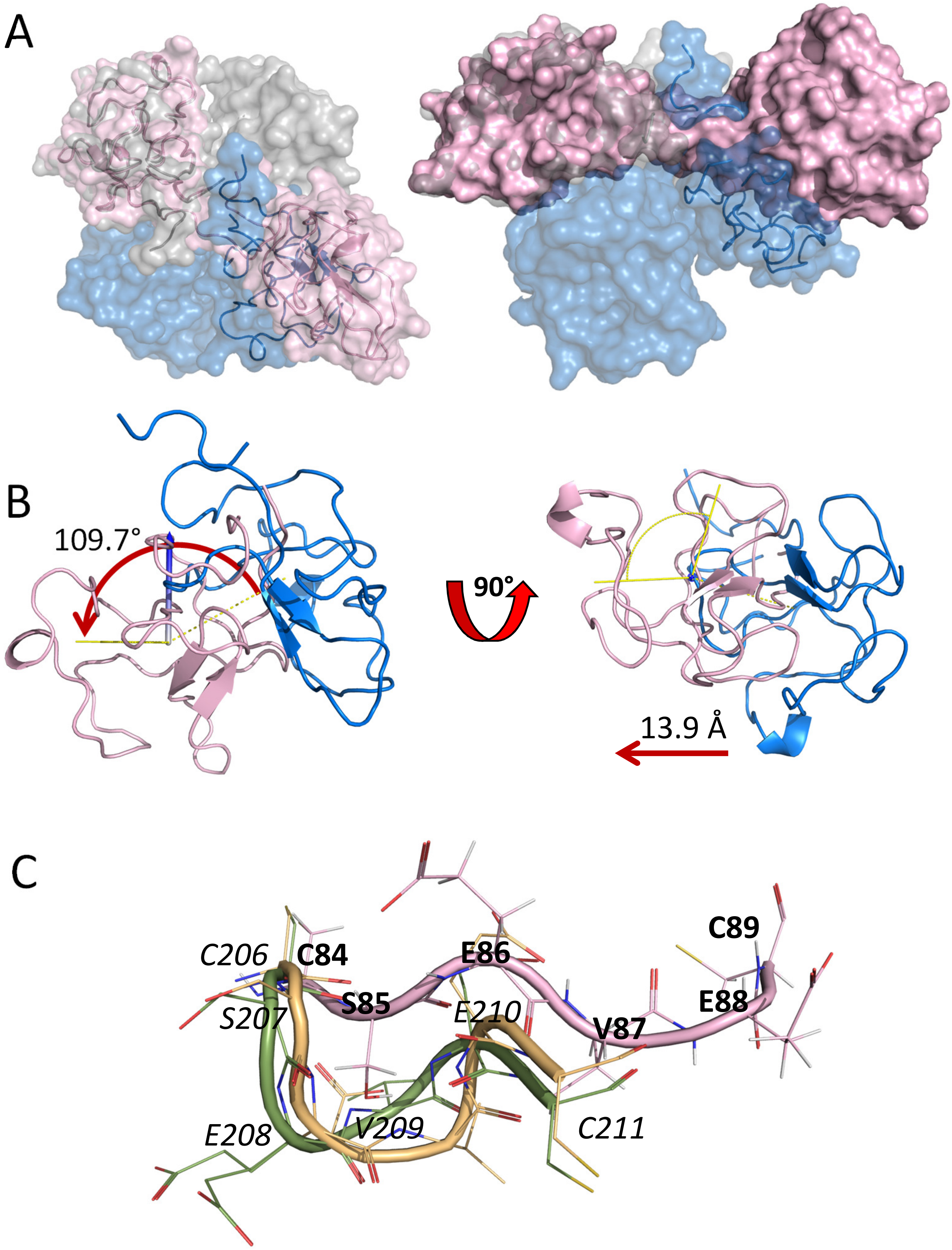

**Supplementary figure S2**

D

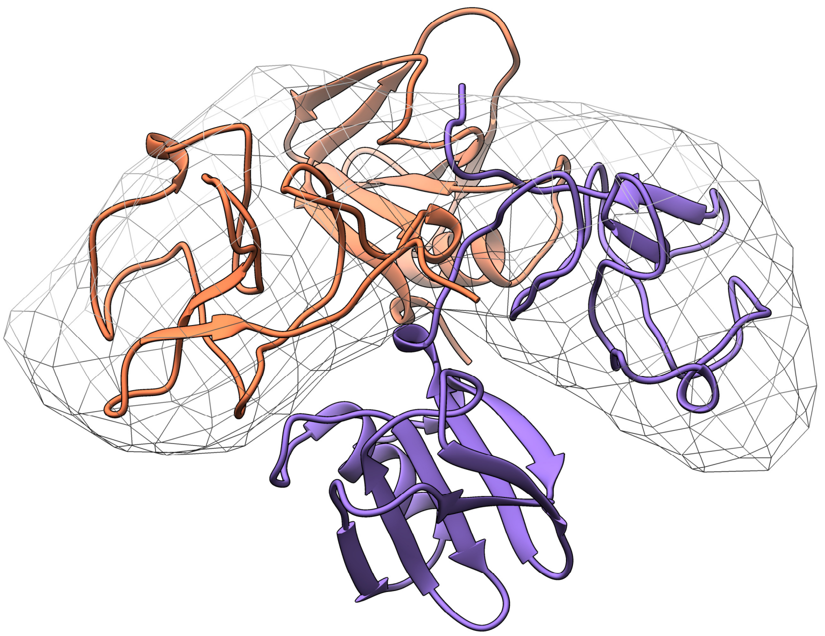

E

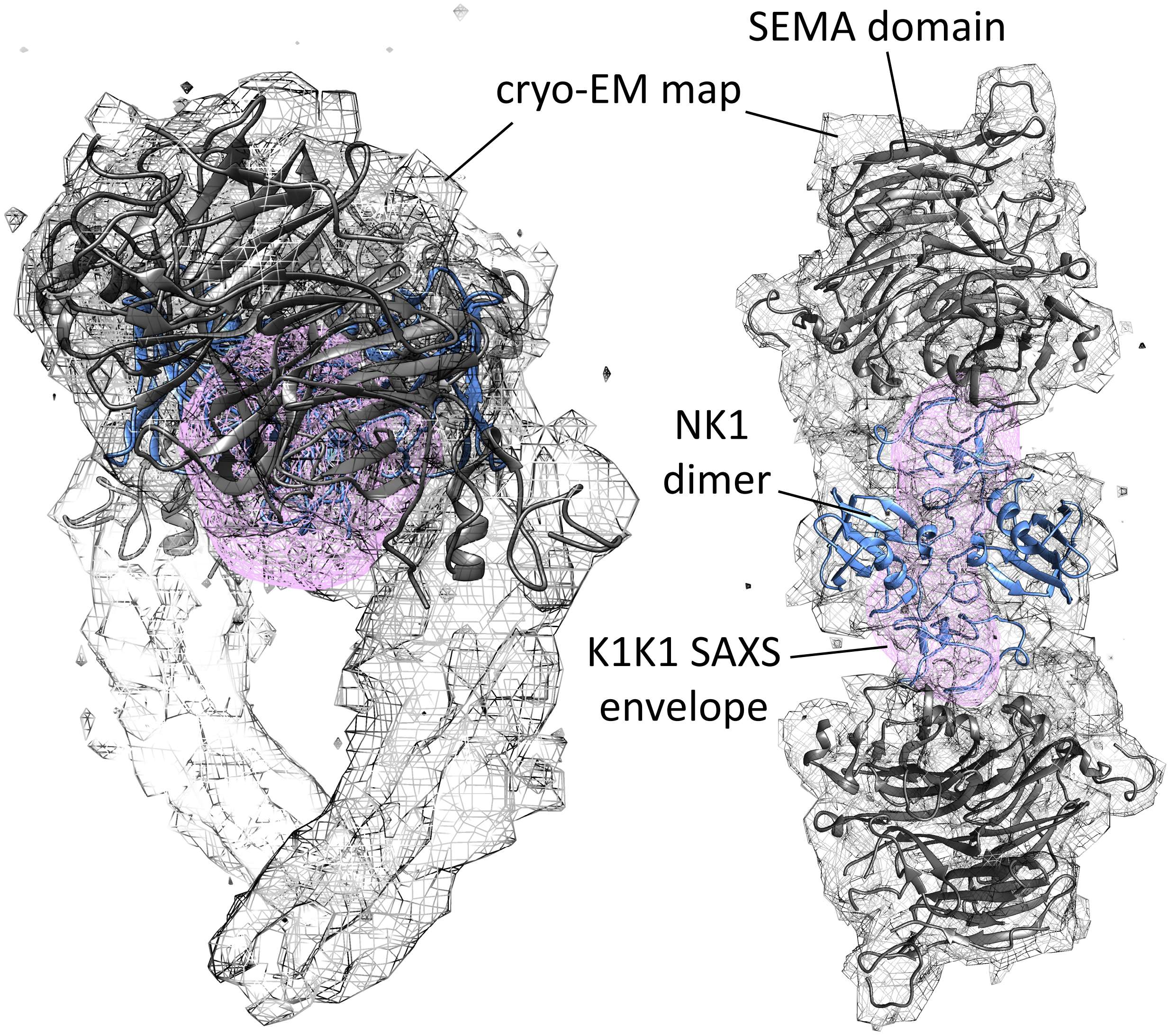

**Supplementary figure S3**

A

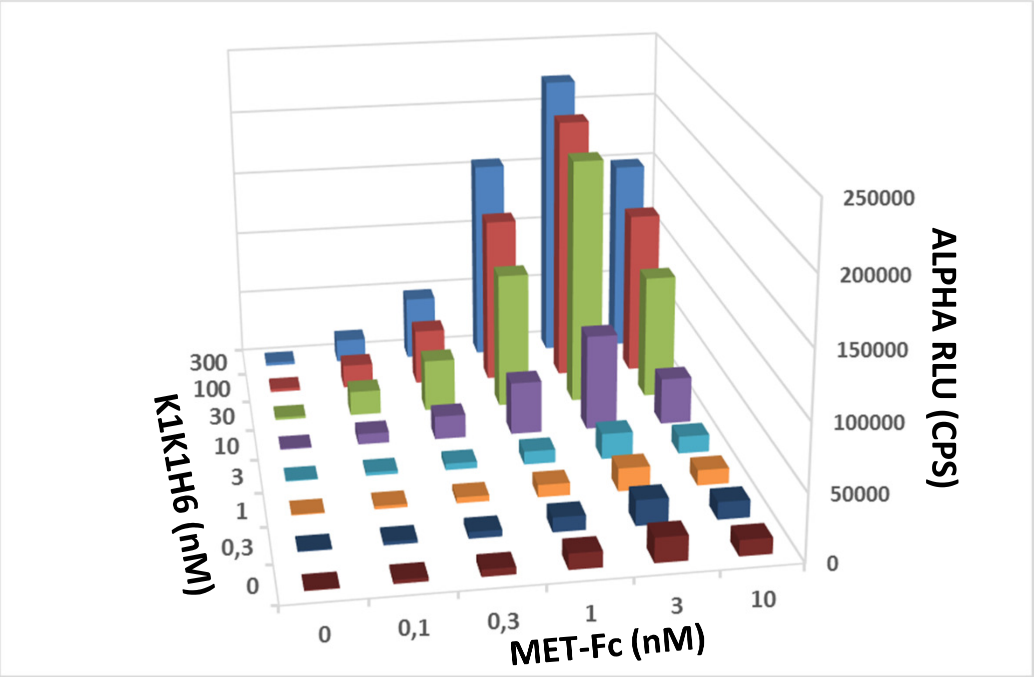


B

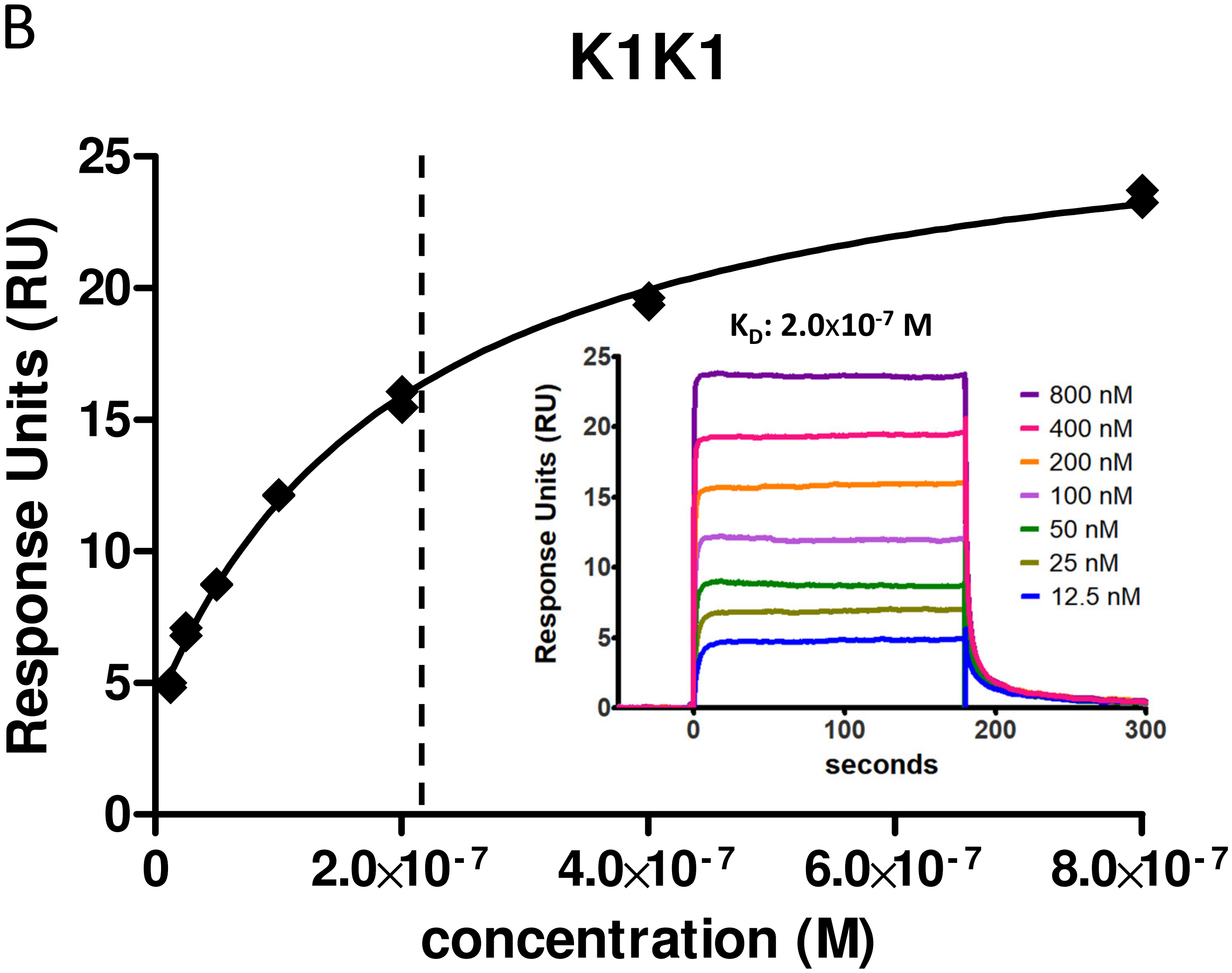

**Supplementary figure S3**

C

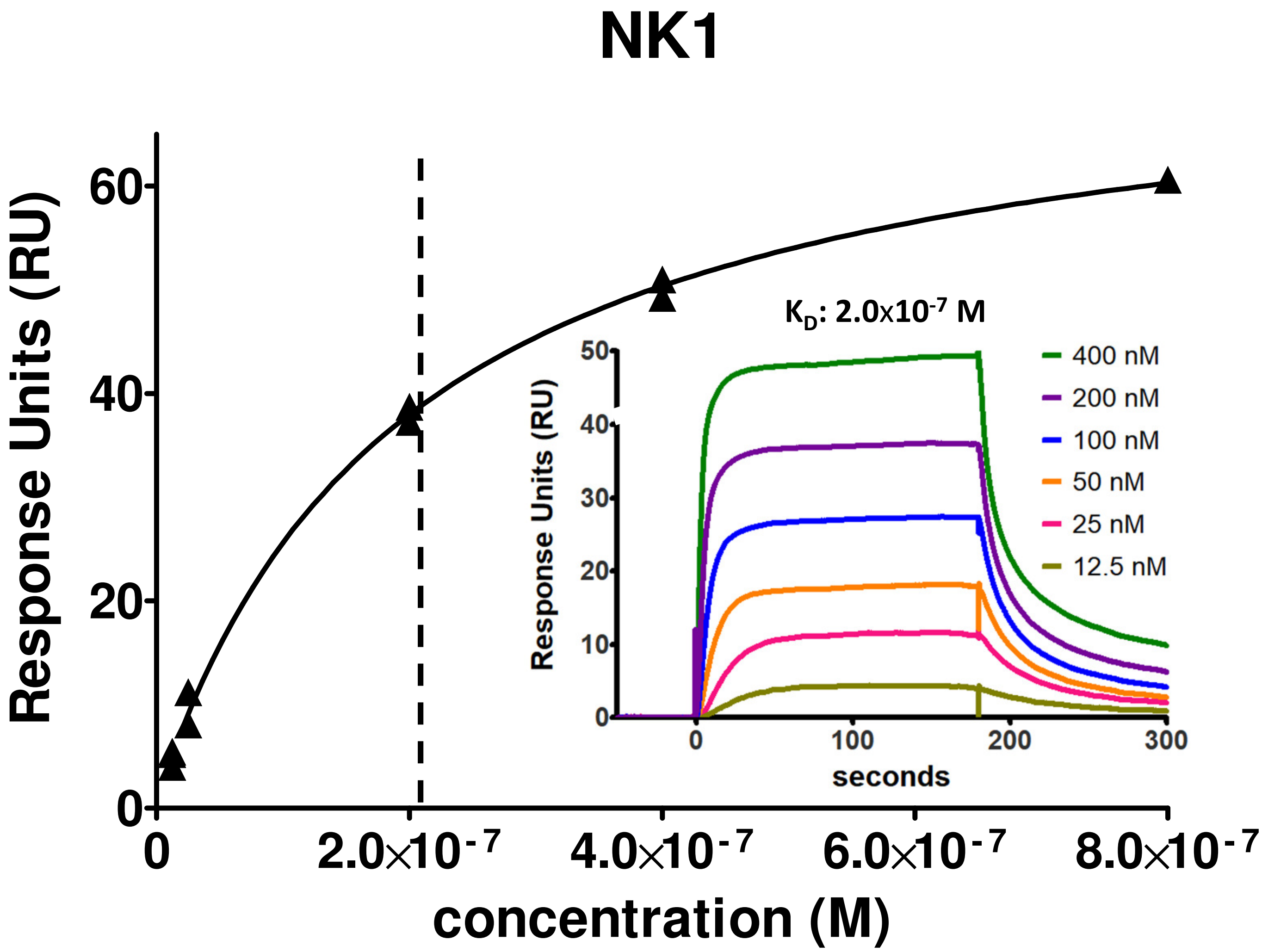

### Supplementary figure S4

A

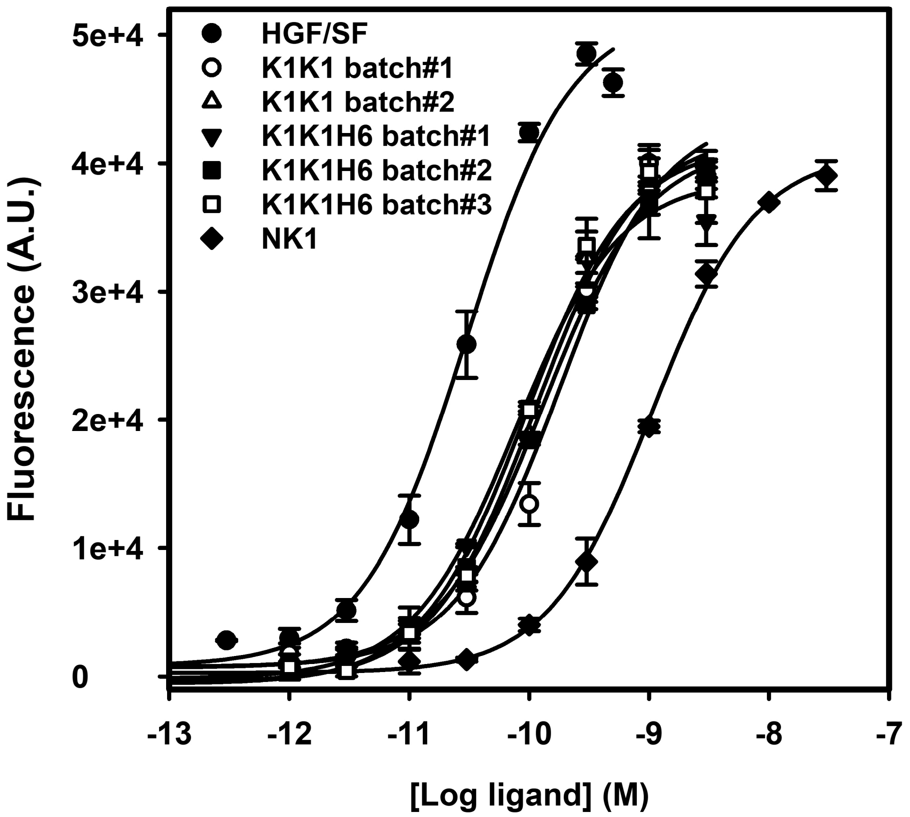


B

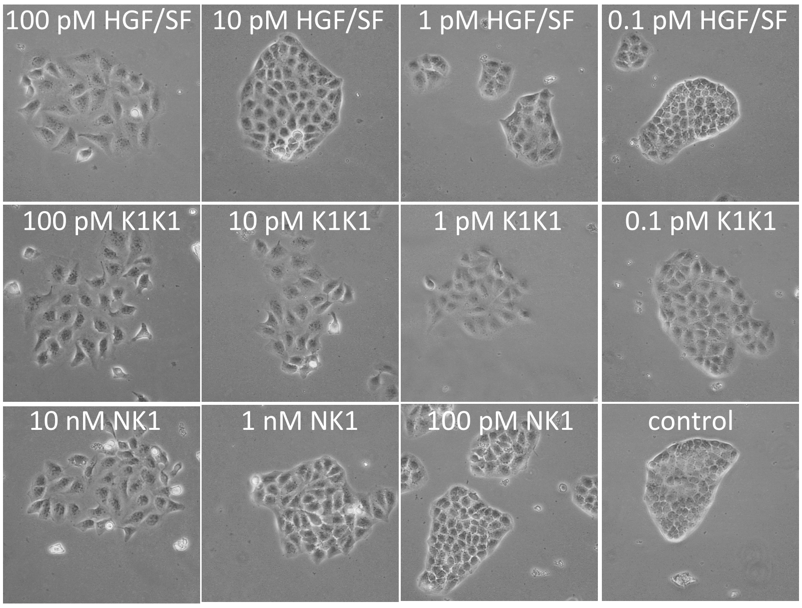

### Supplementary figure S5

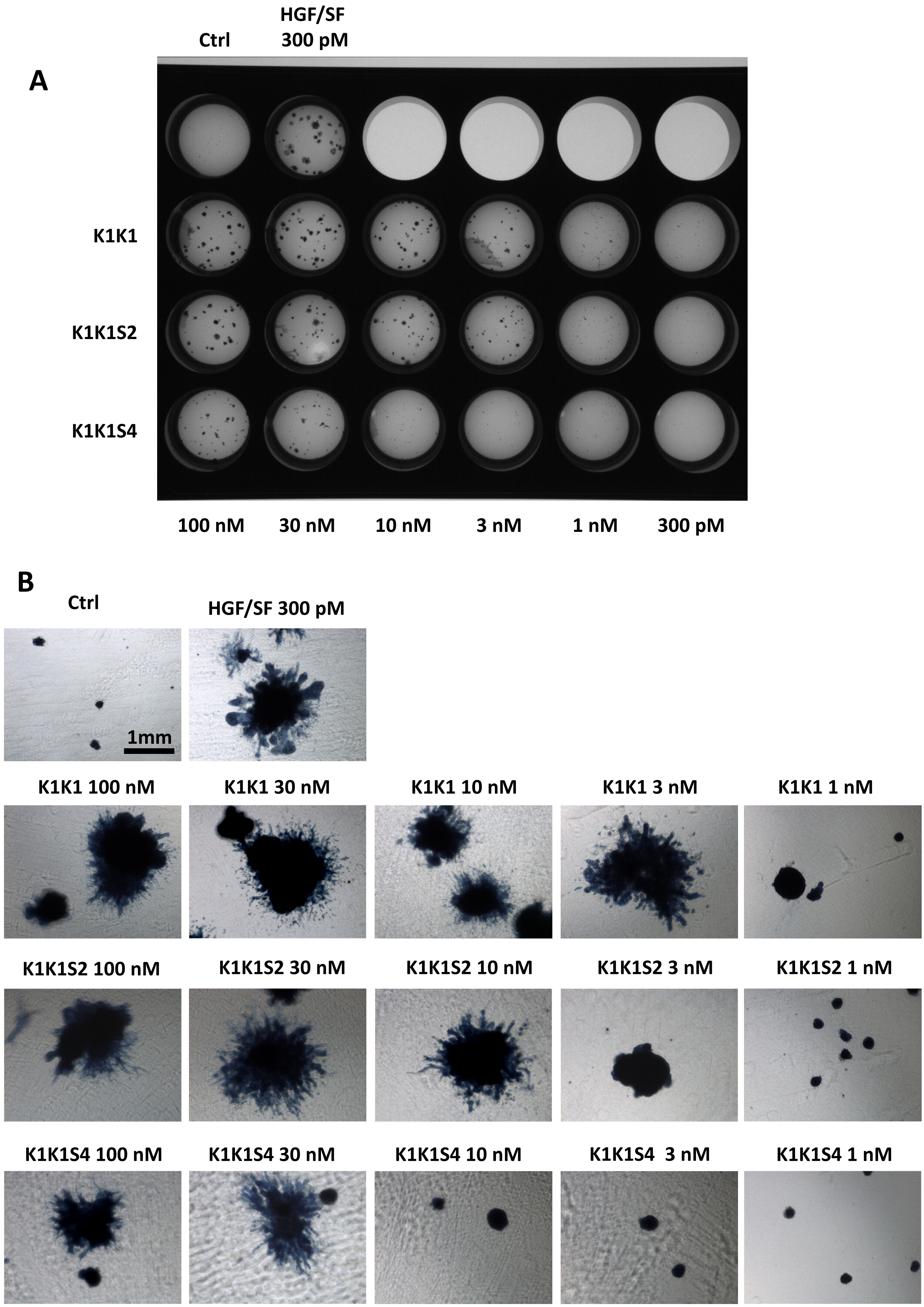

### Supplementary figure S5

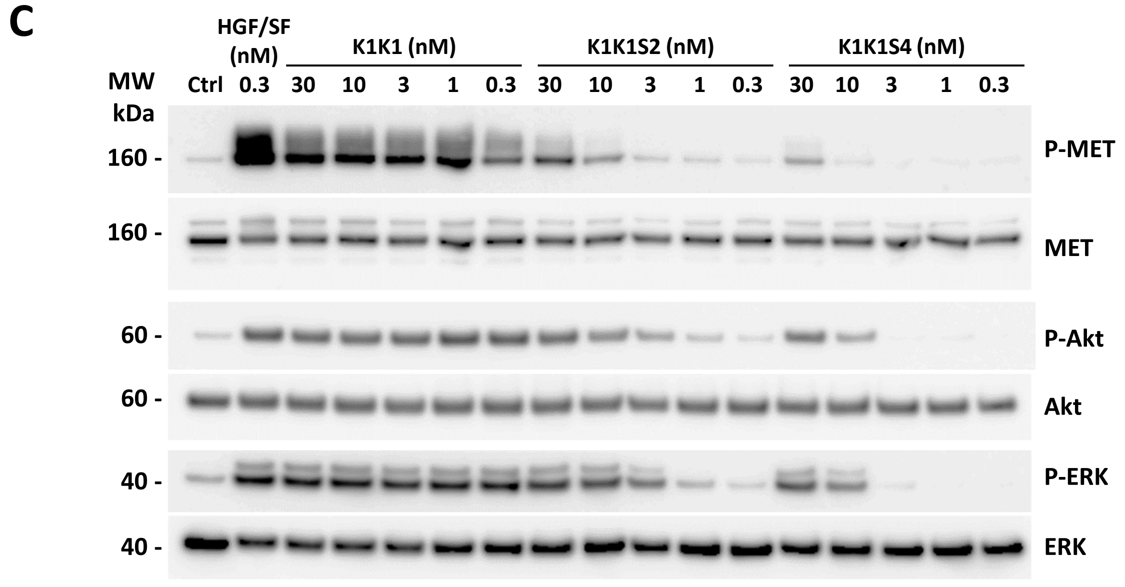

### Supplementary figure S6

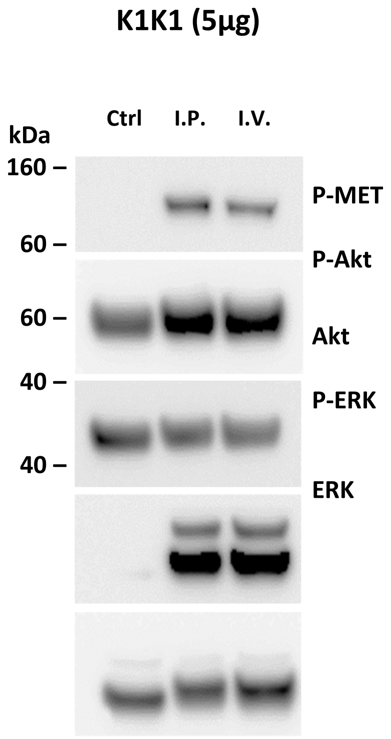

### Supplementary figure S7

## A

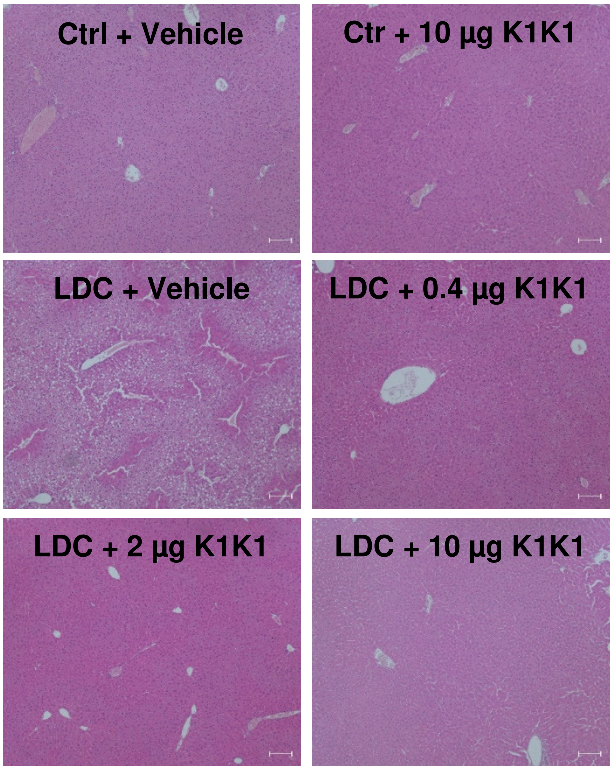


## B

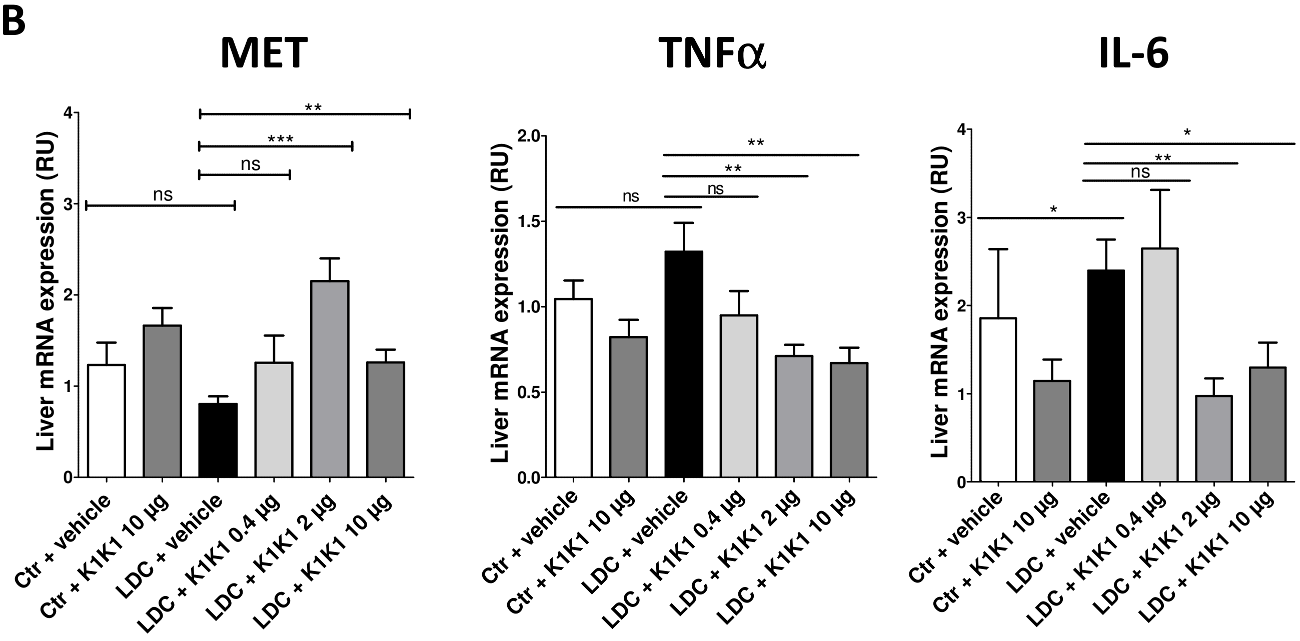

**Supplementary figure S8**

**A**

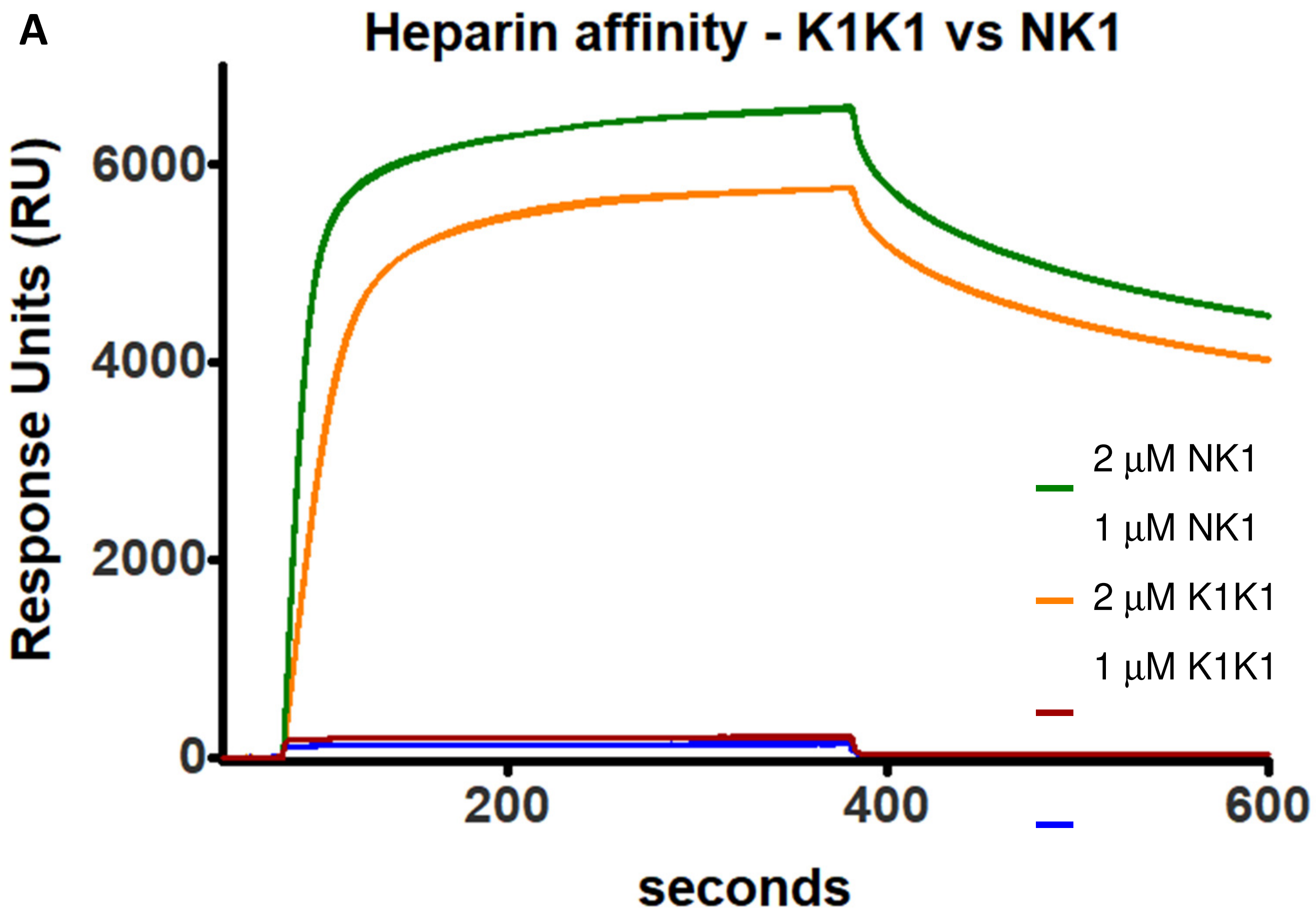


**B**

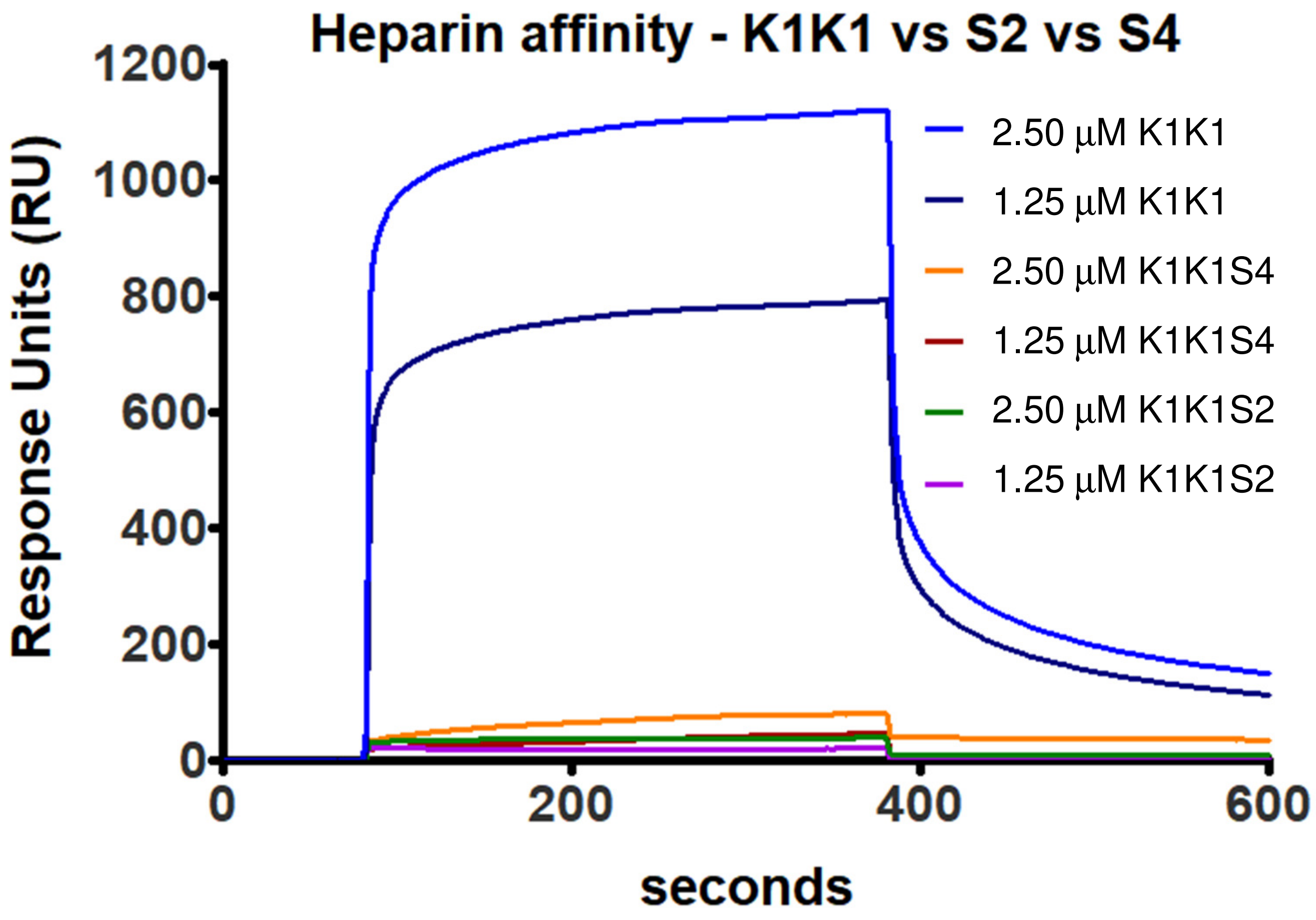

**Supplementary Table S1. Summary of X-ray data collection and refinement statistics.**

| | K1K1H6 | K1K1 |
|---|---|---|
| **Data Collection** | | |
| BeamLine | ESRF ID23-1 | ESRF BM14 |
| Wavelength | 0.9763 | 0.953725 |
| Resolution range (Å) | 35.89 - 1.7 (1.761 - 1.7) | 43.29 - 1.8 (1.864 - 1.8) |
| Space group | P1 21 1 | P1 21 1 |
| Unit cell a, b, c (Å) | 31.8, 59.6, 46.3 90, | 45.57, 58.75, 65.94 |
| a, b, g (°) | 103.8, 90 | 90, 108.219, 90 |
| Total reflections | 69988 (6427) | 161096 (16039) |
| Unique reflections | 18549 (1864) | 30781 (3066) |
| Completeness (%) | 99.81 (99.84) | 99.66 (99.64) |
| Multiplicity | 3.8 (3.4) | 5.2 (5.2) |
| $CC_{1/2}$ | 0.999 (0.971) | 0.986 (0.824) |
| **Refinement** | | |
| $R_{work}$ ($R_{free}$) | 0.1651 (0.1942) | 0.1966 (0.2268) |
| No. of non-hydrogen atoms | 1572 | 2912 |
| Wilson B-factor | 15.16 | 20.27 |
| Average B-factor | 22.03 | 30.04 |
| RMS (bonds) | 0.020 | 0.003 |
| RMS (angles) | 1.68 | 0.62 |
| Ramachandran favored (%) | 97.09 | 95.78 |
| Ramachandran allowed (%) | 2.91 | 4.22 |
| Ramachandran outliers (%) | 0.00 | 0.00 |

*N.B. Statistics for the highest-resolution shell are shown in parentheses.*

**Supplementary Table S2. Summary of SAXS data analysis.**

| | K1K1 | MET567 | K1K1+MET567 |
|---|---|---|---|
| **Data Collection** | | | |
| BeamLine | ESRF BM29 | ESRF BM29 | ESRF BM29 |
| Beam energy (keV) | 12.5 | 12.5 | 12.5 |
| Sample-detector distance (m) | 2.867 | 2.867 | 2.867 |
| Exposure time (s) | 1 | 1 | 1 |
| Sample cell thickness (mm) | 1 | 1 | 1 |
| Sample concentration (mg/mL) | 8.5 mg/mL | 7.6 mg/mL | 12.4 mg/mL |
| Temperature (°C) | 20 | 20 | 20 |
| Final q range ($nm^{-1}$) | 0.01 - 4 | 0.01 - 4 | 0.01 - 4 |
| **Data Analysis** | | | |
| Points used for Guinier analysis | 1-94 | 11-48 | 2-28 |
| Guinier $qR_g$ limits | 1.30 | 0.97 | 0.99 |
| Guinier $R_g$ (nm) | 2.22 | 3.23 | 3.78 |
| I(0) ($mm^{-1}$) | 14.8 ± 0.01 | 64.9 ± 0.04 | 69.9 ± 0.08 |
| $D_{max}$ (nm) | 6.6 | 11.5 | 14.2 |
| MW estimation ($V_c$ based) (kDa) | 16.5 | 66.2 | 83.4 |
