## Supplemental legends and tables for "Dimerization of kringle 1 domain from hepatocyte growth factor/scatter factor provides a potent minimal MET receptor agonist"

### Supplementary figure legends:

**Figure S1. Purity of K1K1 after heparin Sepharose affinity purification.**

(A) Chromatography profile of the elution of K1K1 from Heparin HiTrap™ column showing the 280 nm absorbance in blue and the gradient of eluent (1 M NaCl) in green. (B) UPLC/MS analysis of K1K1 after size exclusion chromatography showing estimated molecular mass nearly identical to the mass as predicted based on the amino acid sequence (19195.24 vs. 19207.69 Da). (C) UPLC/MS analysis of K1K1H6 after size exclusion chromatography and showing estimated molecular mass nearly identical to the mass as predicted based on the amino acid sequence (20018.24 vs. 20030.53 Da). (D) Coomassie stained SDS-PAGE gel showing the different recombinant proteins used in this study (see design in Fig .1). About 5 µg of protein were loaded in reducing sample buffer in each lane.

**Figure S2. Structural alignment of K1K1 with NKI dimer.**

(A) Semi-transparent surface presentation of K1K1 alignment based on one kringle domain with that of the kringle domain of one NK1 protomer. The alignment projects the second K1K1 kringle across and positions it close to but not in the same position as the second kringle domain in the NK1 dimer (blue). (B) Two detailed 90° views of the K1K1-NK1 kringle alignment showing the kringle of K1K1 in pink and the kringle of the NK1 protomer in magenta. The straight, stretched out conformation of K1K1 misaligns the second kringle domain by a rotation of 109.7° leading to a translation of 13.9 Å. (C) The linker based on the naturally occurring linker sequence between kringle 1 and kringle 2 in HGF/SF, SEVE, is straight in K1K1 (pink) and has a different conformation in two NK2 structures available (3HN4 in yellow and 3SP8 in green). Numbering is given in italic from the last cysteine of kringle 1 (C206 in NK2) until the first cysteine in kringle 2 (C211 in NK2) with equivalent K1K1 residues in bold. (D) Superimposition of K1K1 SAXS envelop with the two kringle domains of the protomers in the NK1 dimer. (E) Additional side and top views of the K1K1 SAXS envelope (pink) placed within the cryo-EM map of the NK1 dimer (blue)-SEMA domain (grey) complex (7mob.pdb).

**Figure S3. Characterization of K1K1H6 and K1K1 to MET by AlphaScreen™ and SPR analysis.**

(A) AlphaScreen™ cross titration assay using MET-Fc chimera (0 -10 nM) and K1K1H6 (0 – 300 nM) dilutions was performed using Protein A coated ALPHA acceptor beads with Ni-NTA coated Alpha donor beads. The Alpha signal is expressed in CPS (photon counts per seconds). Measurements are expressed as technical duplicates (mean+/- SD, n=2). (B) SPR analysis of K1K1 to immobilized MET receptor extracellular domain. The main plot shows the binding isotherm with equilibrium response plotted at different concentrations. The inserted graph shows the binding curves at different concentrations of K1K1. (C) SPR analysis of NK1 to immobilized MET receptor extracellular domain. As in (B), the main plot shows the binding isotherm and the inserted graph shows the binding curves at different concentrations of NK1.

**Figure S4: Reproducibility of K1K1 and K1K1H6 batches and efficacy of injection route in mice.**

(A) MDCK cells were treated with 0.7 mM anisomysin for 16 hours with the addition of HGF/SF, NK1, or different batches of K1K1 or K1K1H6 at different concentrations. Plotted are two different batches of K1K1 and three different batches of K1K1H6. Measurements are expressed in raw fluorescence signal as technical duplicates (mean+/- SD, n=2). (B) MDCK cell scattering at different concentrations of K1K1, K1K1S2 and K1K1S3 showing the lowest concentration at which each protein is still active and the subsequent dilution at which no more scattering is observed. HGF/SF and K1K1 1nM were used as positive control (maximum scattering) and PBS as negative control. For complete half-log dilution of agonist series, see source data.

**Figure S5: MDCK cell morphogenesis assay and signalling pathway activation using K1K1, K1K1S2 and K1K1S4 mutants.**

(A) MDCK cell (~500) were seeded into a thick layer of a 1:1 collagen/Growth Factor Reduced Matrigel™ and treated with semi-log dilutions (100 pM to 300 nM) of HGF/SF, K1K1, K1K1S2 and K1K1S4 twice a week for one month. The cells were fixed and stained with Evans Blue and DAPI. Image of the full plate is presented. (B) Colonies were observed in bright field using inverted microscope observed 4x objective on a Nikon Eclipse TS100 microscope. (C) Phosphorylation analysis of MET signalling pathway by Western blot on HeLa cell lysates after stimulation with semi-log dilution (0.3 to 30 nM) of K1K1, K1K1S2 or K1KS4 for 10 min. Loading controls are based on total MET, total Akt, and total ERK present in each lane.

**Figure S6: In vivo MET activation.**

8-week old FVB mice were injected with PBS (Ctrl) or 5 µg of K1K1 either per intravenous (I.V.) or intraperitoneal (I.P.) route of injection. Mice were sacrificed 10 minutes after injection after which MET, Akt, and ERK phosphorylation in liver homogenate was determined by Western blot. Blot present total ERK and Akt proteins as loading controls.

**Figure S7. *In vivo* evaluation of the efficacy of K1K1 in a mouse model of alcoholic steatohepatitis.**

(A) Hematoxylin-erythrosin B staining of mouse livers submitted to an adapted Lieber DeCarli model (x10). Analyses were performed on 10 animals in control groups (Ctrl + Vehicle and Ctrl +10 µg K1K1) and 15 animals in ethanol treated groups (LDC ± K1K1). Scale bar = 100 µm. (B) The mRNA expression of MET and inflammatory cytokines TNFα and IL-6 was analysed in mouse livers using RT-qPCR with b-actin as housekeeping gene. Results are expressed relative unit (RU) and represented as mean ± SD.

**Figure S8: SPR analysis of heparin binding by NK1, K1K1, K1K1S2, and K1K1S4.**

(A) NK1 and K1K1 were each injected at 2 μM and 1 μM on a heparin-coated SC HEP0320.a chip (Xantec). (B) Comparison of heparin affinity of K1K1, K1K1S2, and K1K1S4, injected at 2.5 μM and 1.25 μM each. All injections were done with a flow of 30 μl/min at 25 °C in PBS running buffer.
