## Supplementary material for "Dimerization of kringle 1 domain from hepatocyte growth factor/scatter factor provides a potent minimal MET receptor agonist": image source data

### Images Data Source File

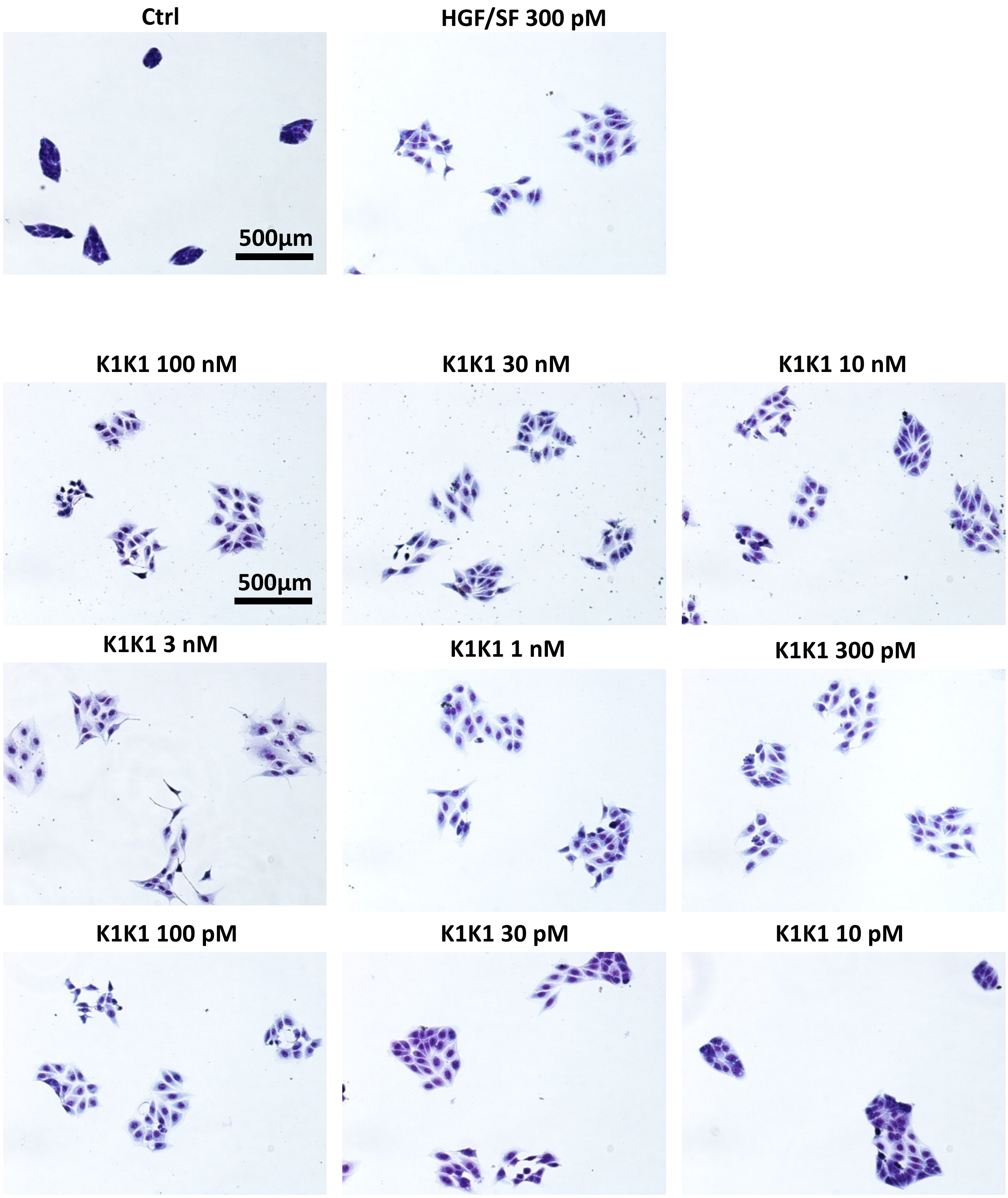


X40 Nikon Eclypse Bright Field

### Images Data source Files

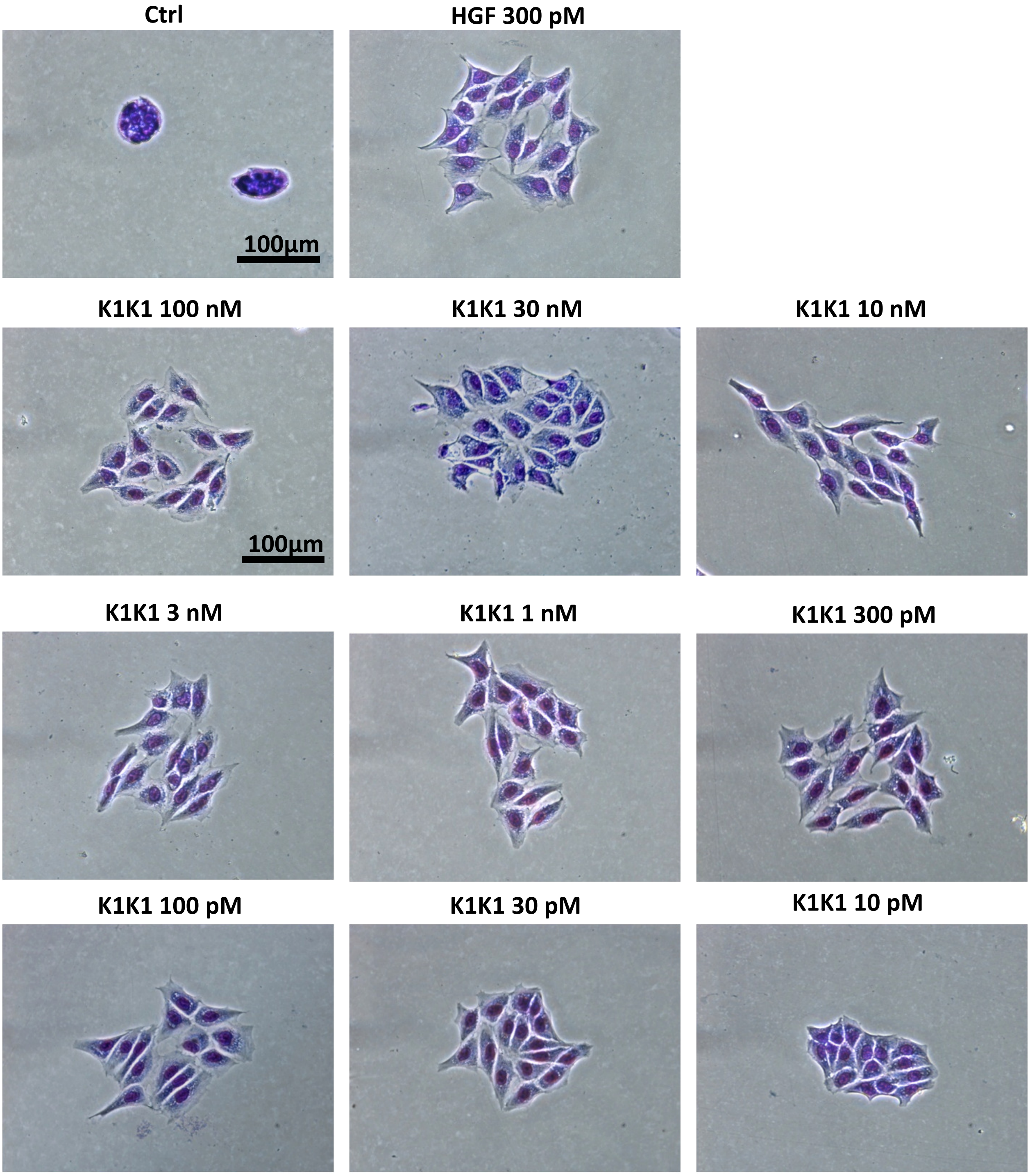


X200 Nikon Eclypse Bright Field

### Data Source File

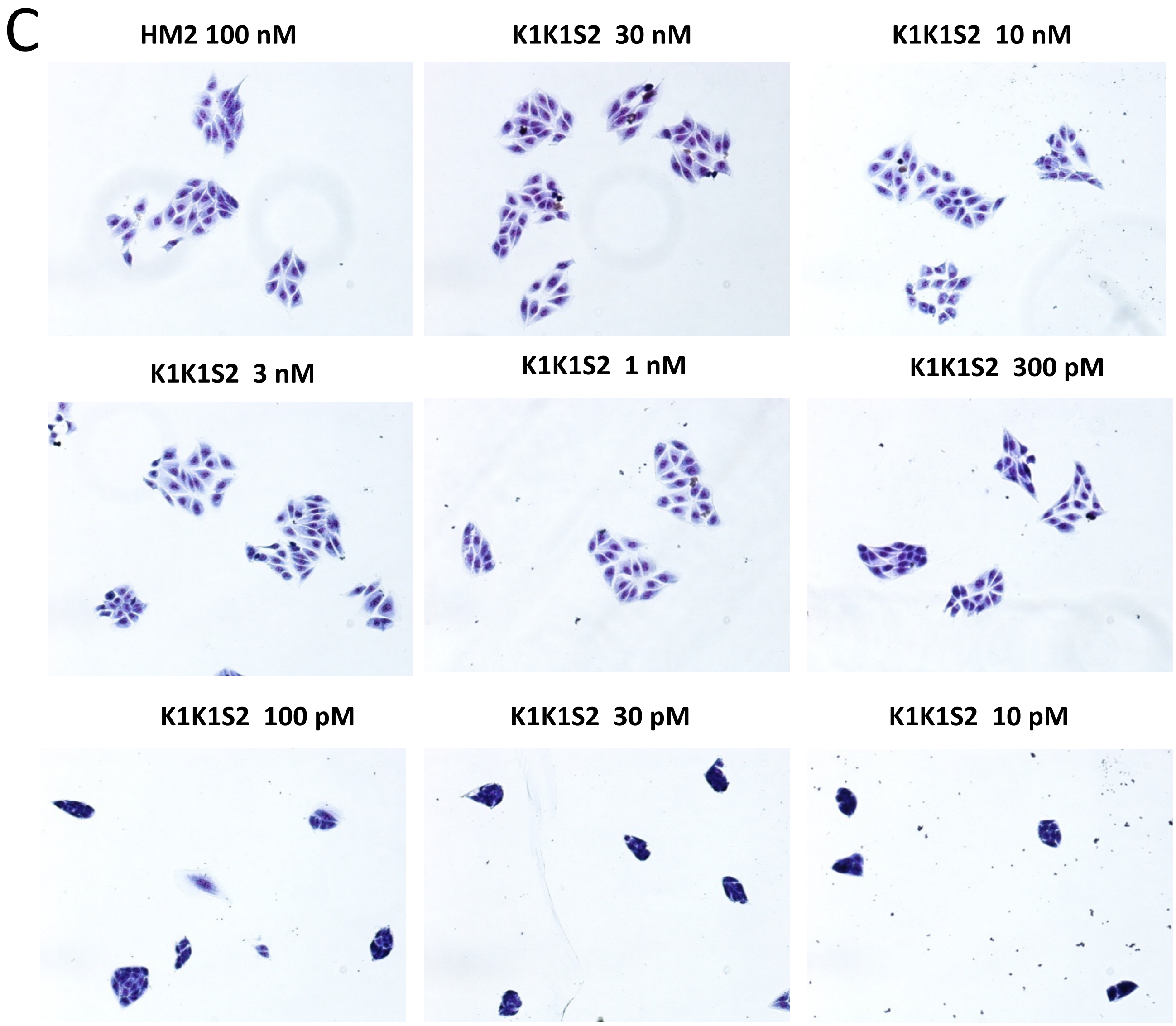


X40 Nikon Eclypse Bright Field

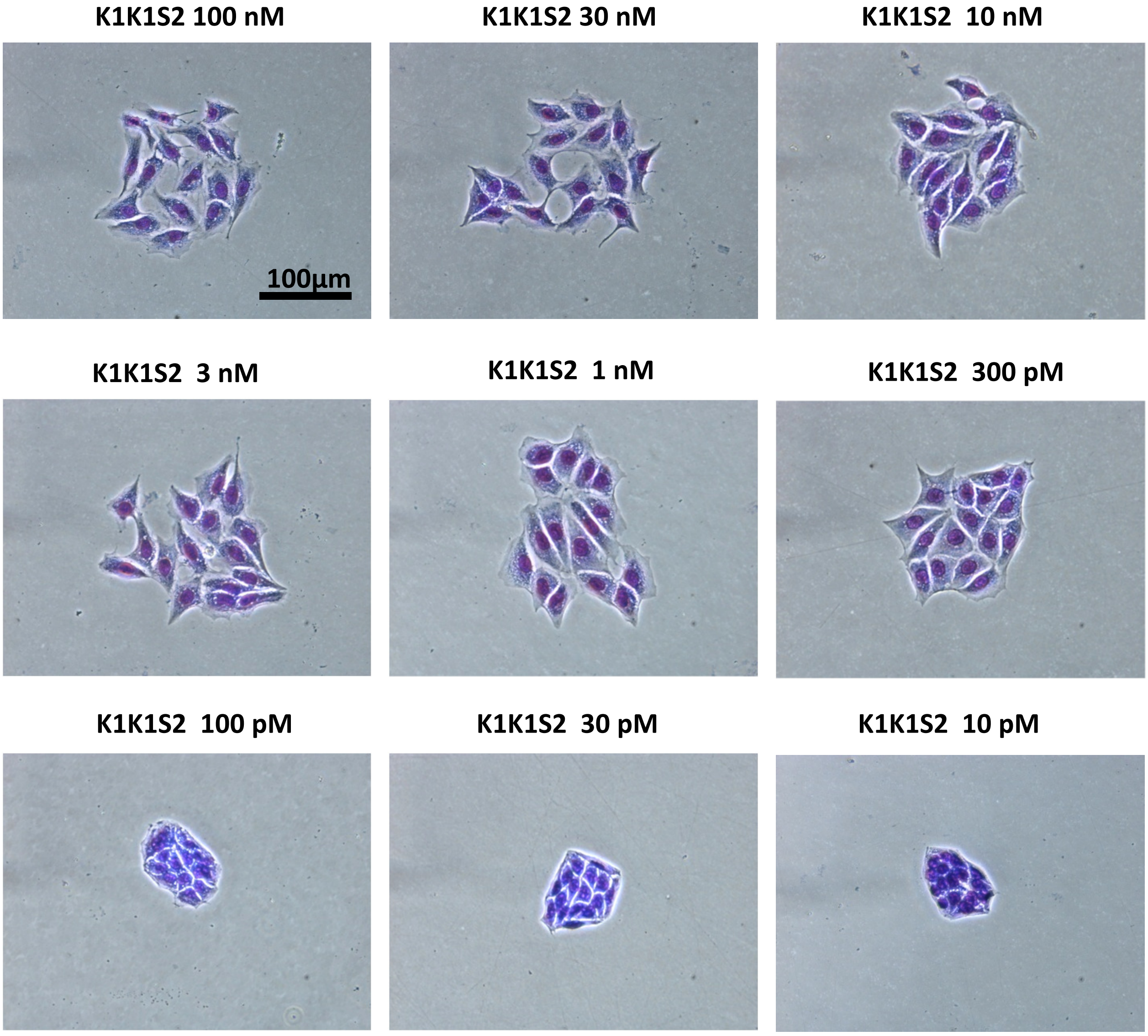


X200 Nikon Eclypse Bright Field

**Data Source File**

D

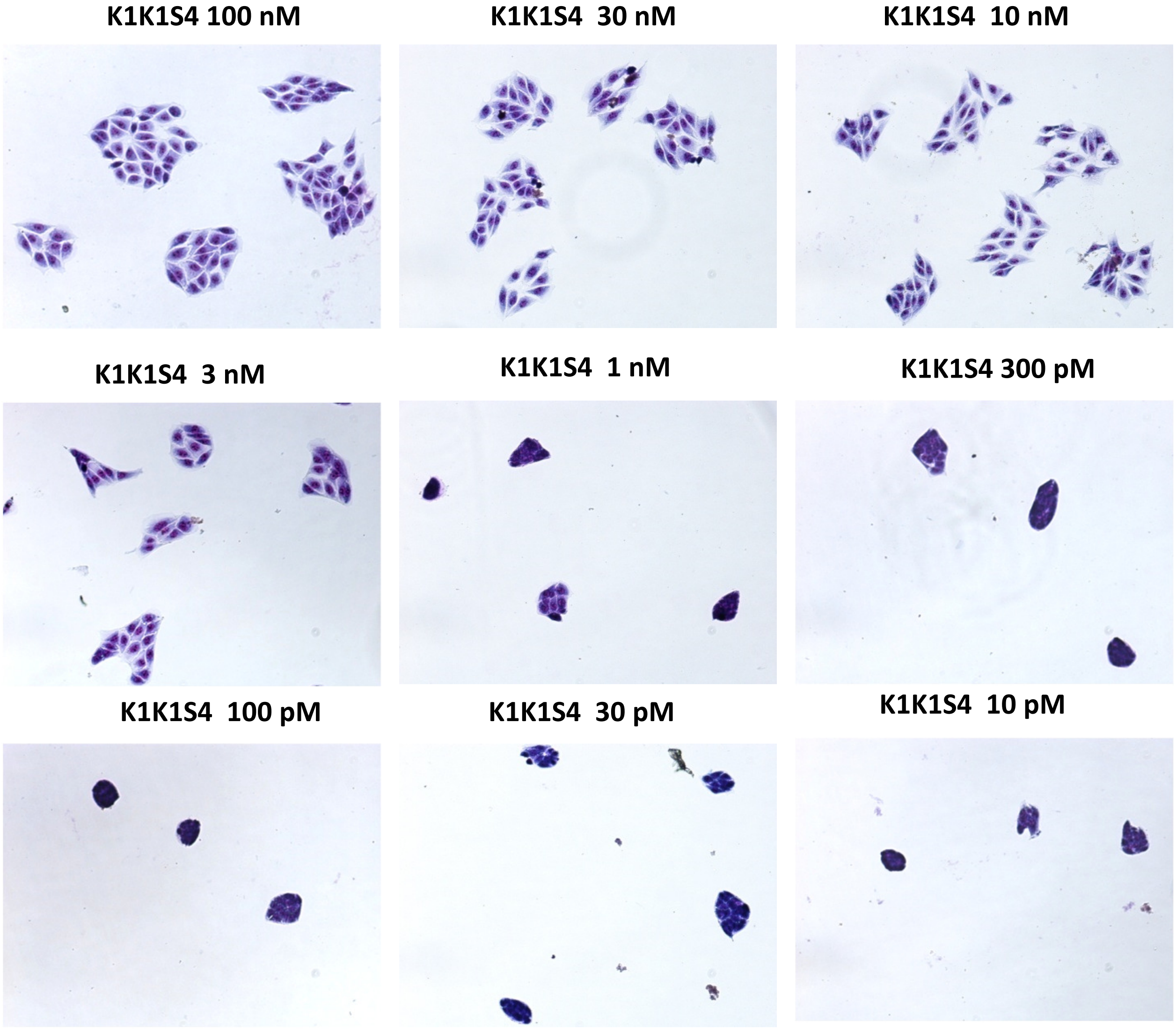


X40 Nikon Eclypse Bright Field

### Data Source File

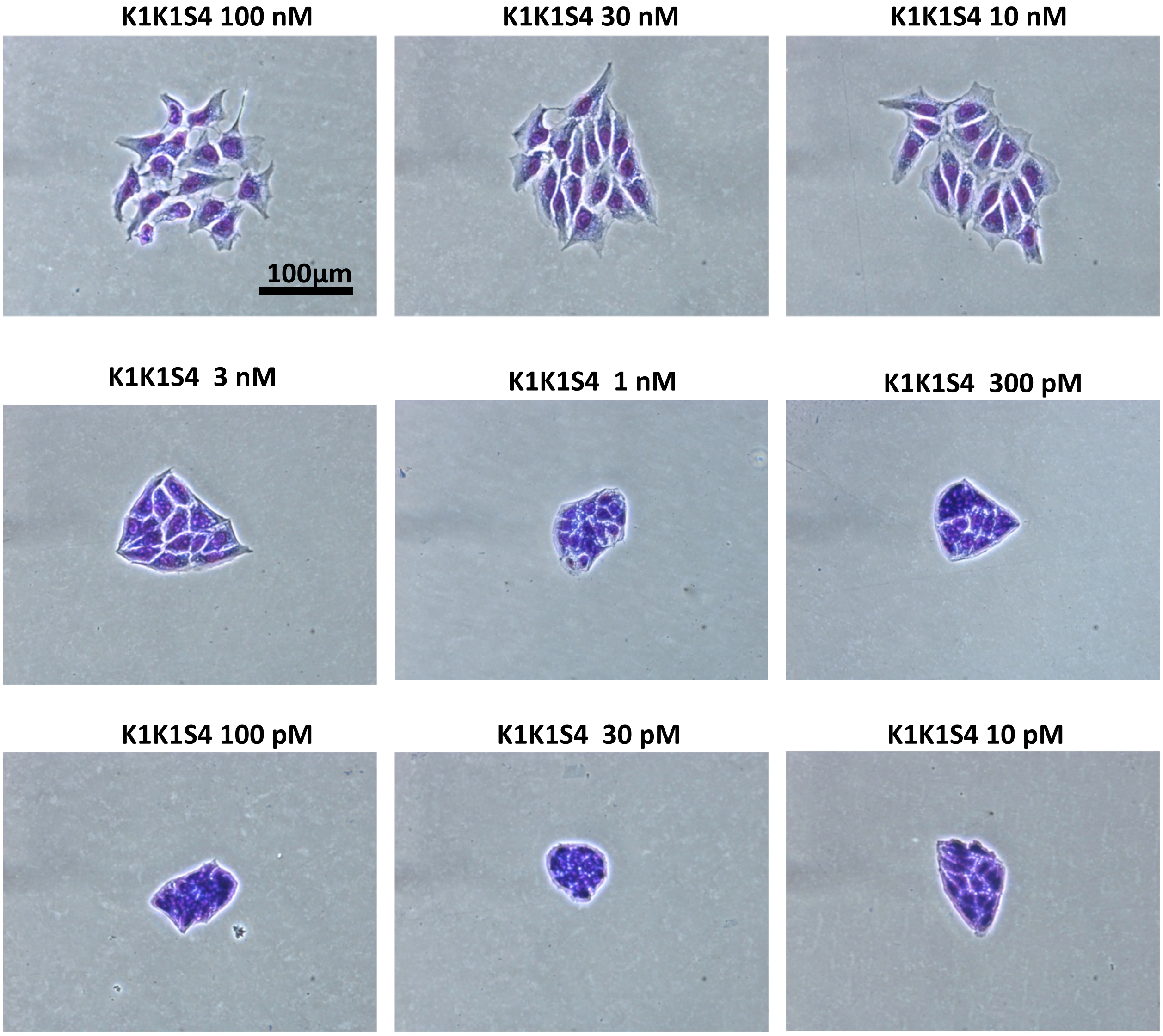


X200 Nikon Eclypse Bright Field
